# Edaravone activates the GDNF/RET neurotrophic signaling pathway and protects mRNA-induced motor neurons from iPS cells

**DOI:** 10.1101/2021.05.09.443325

**Authors:** Qian Li, Yi Feng, Yingchao Xue, Xiping Zhan, Yi Fu, Gege Gui, Weiqiang Zhou, Jean-Philippe Richard, Arens Taga, Pan Li, Xiaobo Mao, Nicholas J. Maragakis, Mingyao Ying

## Abstract

**Background:** Spinal cord motor neurons (MNs) from human iPS cells (iPSCs) have wide applications in disease modeling and therapeutic development for amyotrophic lateral sclerosis (ALS) and other MN-associated neurodegenerative diseases. We need highly efficient MN differentiation strategies for generating iPSC-derived disease models that closely recapitulate the genetic and phenotypic complexity of ALS. An important application of these models is to understand molecular mechanisms of action of FDA-approved ALS drugs that only show modest clinical efficacy. Novel mechanistic insights will help us design optimal therapeutic strategies together with predictive biomarkers to achieve better efficacy.

**Methods:** We induce efficient MN differentiation from iPSCs in 4 days using synthetic mRNAs coding two transcription factors (Ngn2 and Olig2) with phosphosite modification. These MNs after extensive characterization were applied in electrophysiological and neurotoxicity assays as well as transcriptomic analysis, to study the neuroprotective effect and molecular mechanisms of edaravone, an FDA-approved drug for ALS, for improving its clinical efficacy.

**Results:** We generate highly pure and functional mRNA-induced MNs (miMNs) from normal and ALS iPSCs, as well as embryonic stem cells. Edaravone alleviates H_2_O_2_-induced neurotoxicity and electrophysiological dysfunction in miMNs, demonstrating its neuroprotective effect that was also found in the glutamate-induced miMN neurotoxicity model. Guided by the transcriptomic analysis, we show a previously unrecognized effect of edaravone to induce the GDNF receptor RET and the GDNF/RET neurotrophic signaling *in vitro* and *in vivo*, suggesting a clinically translatable strategy to activate this key neuroprotective signaling. Notably, edaravone can replace required neurotrophic factors (BDNF and GDNF) to support long-term miMN survival and maturation, further supporting the neurotrophic function of edaravone-activated signaling. Furthermore, we show that edaravone and GDNF combined treatment more effectively protects miMNs from H_2_O_2_-induced neurotoxicity than single treatment, suggesting a potential combination strategy for ALS treatment.

**Conclusions:** This study provides methodology to facilitate iPSC differentiation and disease modeling. Our discoveries will facilitate the development of optimal edaravone-based therapies for ALS and potentially other neurodegenerative diseases.

**Graphic Abstract:** 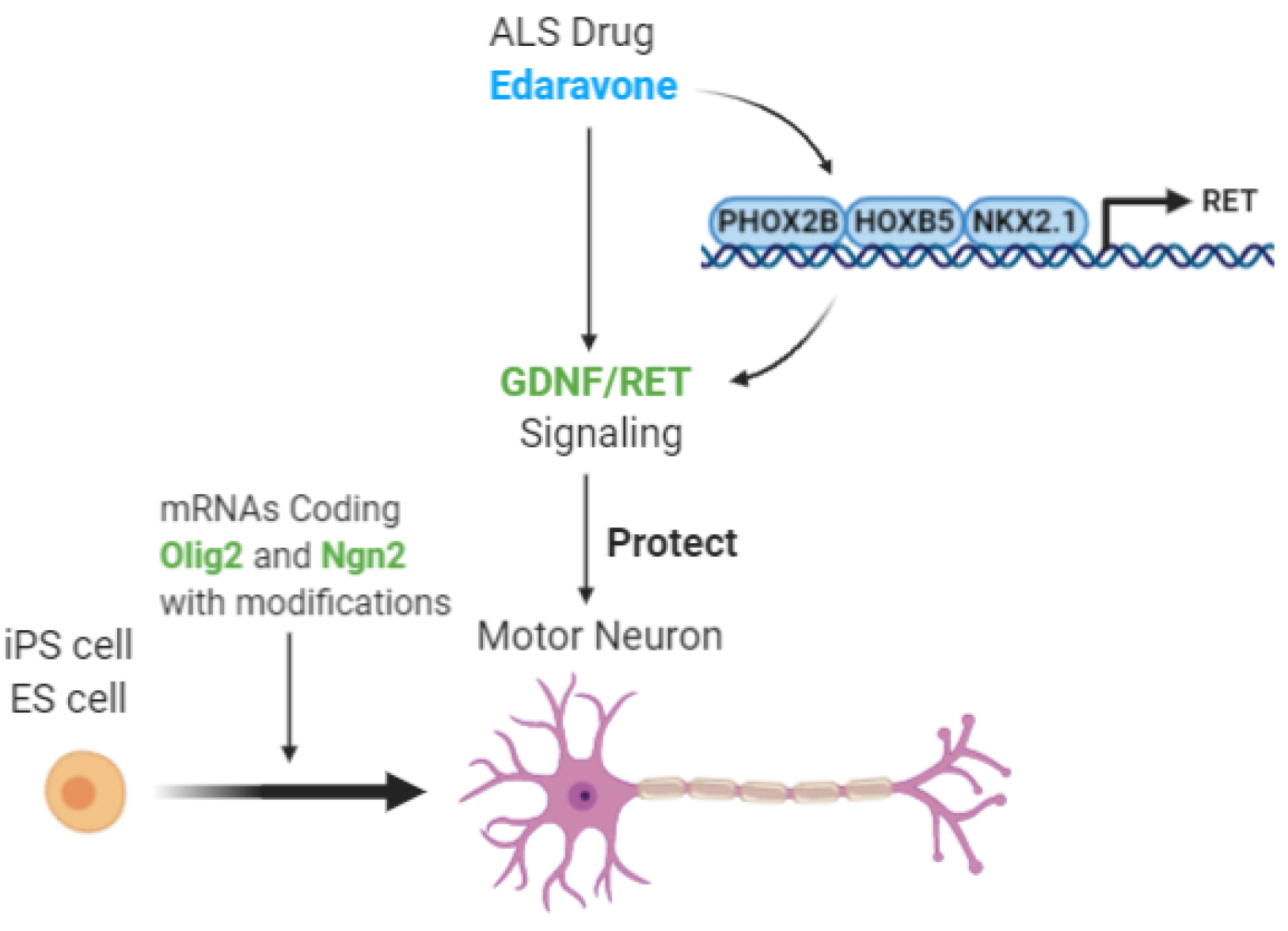

## Background

Human pluripotent stem cells (PSCs), including embryonic stem cells (ESCs) and induced pluripotent stem cells (iPSCs), have unique characteristics, such as long-term self-renewal and multi-lineage differentiation capability. Patient-derived iPSCs are widely used in regenerative medicine to provide disease-relevant functional cells for mechanistic studies, drug discovery and cell replacement therapy [1–3]. All iPSC-based applications rely on robust differentiation strategies for the manufacture of lineage-specific and functional progenies.

Human motor neurons (MNs) derived from iPSCs provide a unique and efficient platform for modeling various MN disorders and developing effective therapies. Amyotrophic lateral sclerosis (ALS) is a rapidly progressive neurodegenerative disease that is characterized primarily by MN degeneration in the brain and spinal cord and has a median survival of 20 to 48 months [4]. Therapeutic development for ALS is extremely challenging [5]. The Food and Drug Administration (FDA) has only approved two drugs with modest efficacy. The first FDA-approved ALS drug riluzole is a glutamatergic neurotransmission inhibitor, and only improves patient survival by 2-3 months without showing benefit on motor function [6–8]. The recently approved edaravone (also known as Radicava or MCI-186) was shown to slow early-stage disease progression in a subset of ALS patients enrolled in a phase III study [9]. However, controversy remains over the clinical efficacy of edaravone in ALS patients [10–12]. It is also unclear how edaravone might be effective in ALS, although it is predicted to act through reducing oxidative stress based on its proposed function as a free radical scavenger [13, 14]. To improve clinical efficacy of current ALS drugs, it is critical to understand their molecular mechanisms of action in disease models that closely recapitulate the genetic and phenotypic complexity of ALS, and further rationally design single or possibly combination therapies together with predictive biomarkers to achieve better clinical efficacy.

ALS is a highly heterogeneous disease with regard to phenotype and progression, a large number of disease-associated genetic variants, and multiple cellular pathways affected [5]. A bank of iPSC-derived MNs from familial and sporadic ALS patients provides an efficient modeling system mimicking the complexity of ALS and suitable for high-throughput drug screening. Building this MN bank needs a robust iPSC differentiation strategy. Traditional MN differentiation methods mainly use small molecule compounds to induce neural conversion and MN lineage specification in iPSCs. These multi-step protocols take about 10-14 days to generate MN precursors with variable purity (commonly 50-70%) [15, 16]. Since the principle of all MN differentiation strategies is based on generating Olig2^+^ MN progenitors [15], it may be feasible to drive robust MN conversion through ectopic expression of Olig2 and other transcription factor (TF) drivers of MN development (e.g. Ngn2) [17, 18]. Ectopic TF expression can be achieved by synthetic mRNA delivery, an efficient, non-viral and non-integrating strategy that has been used by us to differentiate iPSCs to dopaminergic neurons [19].

Here, we established a rapid MN differentiation method using synthetic mRNAs coding two TFs. These MNs were applied to high-throughput phenotypic analysis and transcriptomic profiling to study the ALS drug edaravone. We revealed a novel effect of edaravone in activating the neurotrophic factor signaling pathway *in vitro* and *in vivo*, and found an effective combination strategy to protect MNs.

## Materials and Methods

### Cell Culture

The normal iPSC line N1 and N3 (referred to as iPSC1 and iPSC3) was derived from human skin fibroblasts as previously characterized and used by us [19, 20]. Two ALS iPSC lines (referred to as iPSC2 and iPSC4) was derived from skin fibroblasts from an ALS patient with the *SOD1^A4V^* mutation. Cell reprogramming was performed using the Sendai virus system with SOX2/OCT4/KLF4/MYC (CytoTune-iPS Reprogramming Kit; Thermo Fisher Scientific, Rockville, MD, http://www.thermofisher.com). iPSC pluripotency is characterized by immunocytochemistry for pluripotent markers (NANOG, OCT4, TRA-1-60, and SSEA-3) and embryoid body formation assay. The human ESC line H1 was obtained from WiCell Research Resources (Madison, WI, http://www.wicell.org). iPSCs and ESCs were maintained in mTESR1 medium (Stem Cell Technologies, Vancouver, BC, Canada) at 5% CO_2_/95% air condition at 37°C and were passaged using ReLeSR™ (Stem Cell Technologies). Karyotype analysis of G-banded metaphase chromosomes has been performed to confirm chromosomal integrity. Human neural stem cells were established from a fetal brain, immortalized by *v-myc* and extensively characterized by Dr. Vescovi and his colleagues [21]. Astrocyte differentiation uses DMEM/F12 medium (Thermo Fisher Scientific) with 1% fetal bovine serum (FBS) for 21 days following the publication [21].

All chemicals were from Sigma Aldrich unless otherwise mentioned.

### mRNA Synthesis and Transfection

Coding sequences of human Ngn2 and Olig2 were cloned into a vector containing the T7 promoter and poly(A) tail for in-vitro transcription as reported by us [19]. mRNA transfection used the Lipofectamine™ Stem Transfection Reagent (Thermo Fisher Scientific). For each well of the 12-well plate, we used 0.25 µg mRNAs (N-SA:O-SA = 1:1) with 1.5 µl lipid. All procedures involving recombinant DNA follow the National Institutes of Health guidelines.

### mRNA-induced MN Differentiation

iPSCs were plated at a density of 3 x 10^5^ cells per well in a 12-well plate coated with growth-factor-reduced Matrigel (Corning). iPSCs were transfected daily with N-SA/O-SA mRNAs for 3 days. Culture medium with SHH (100 ng/ml) and DAPT (10 µM) were changed daily and shifted from mTeSR1 to N2 (Thermo Fisher Scientific) in 3 days. Cells were dissociated by Accutase and replated to poly-D-Lysine/Laminin-coated surface at the density of 1×10^5^ cells/cm^2^. Neuron maturation medium contain neurobasal medium with the B27 supplement, BDNF (10 ng/ml), GDNF (10 ng/ml), cAMP (0.1 mM), ascorbic acid (0.2 mM), DAPT (10 µM). Medium was changed after 48h followed by half change every 3-4 days. Cryopreservation medium containing 40% neurobasal medium, 50% FBS and 10% DMSO.

### Western Blotting

We follow our previous publication [19] to perform total protein extraction using RIPA buffer (Sigma-Aldrich), SDS-polyacrylamide gel electrophoresis and western blotting. Protein levels were quantified with the Odyssey IR Imaging System (LI-COR Biosciences). All primary antibodies are listed in Table S6.

### Immunofluorescence staining and quantification

Cells were fixed in 4% paraformaldehyde in PBS (pH 7.4) and subjected to immunostaining as published by us [19], using primary antibodies listed in Table S6. The percentage of marker-positive cells over DAPI+ nuclei was determined in samples from at least three samples that were independently differentiated for staining. The high-content analysis software (HCA-Vision V2.2.0. CSIRO) was used for nucleus detection and cell body segmentation. Threshold for each marker was set based on signal intensity in the IgG isotype control.

### GDNF ELISA

ELISA used the kit from Thermo Fisher Scientific (human GDNF) and Rockland (mouse GDNF).

### Electrophysiological Recordings

Voltage-clamp recording was performed at 35°C in a chamber perfused with regular artificial cerebrospinal fluid (124 mM NaCl, 2.5 mM KCl, 1.3 mM MgCl_2_, 2.5 mM CaCl_2_, 1mM NaH_2_PO_4_, 26.2 mM NaHCO_3_, 20 mM glucose, pH 7.4, equilibrated with 95% O_2_ and 5% CO_2_, ∼310 mosm), which flowed at 3 ml/minute. Patch electrodes were pulled from borosilicate glass and had resistances of 2.0 – 4.0 MΩ when filled with an intracellular solution (135 mM KMeSO_4_, 5 mM KCl, 5 mM HEPES, 0.25 mM EGTA-free acid, 2 mM Mg-ATP, 0.5 mM GTP, 10 mM phosphocreatine-tris, pH 7.3, ∼290 mosm). Neurons were identified using a 10X objective mounted on an upright microscope with transmitted light, and their neuronal somata were then visualized through a 40X water immersion objective using IR differential interference contrast optics (DIC). The cell somatic recordings were made using an Axopatch 700B amplifier in combination with pClamp 11 software (Molecular Devices). Neurons were initially voltage-clamped at -70 mV, and R_series_ and R_input_ were monitored using a 2.5-mV 100-ms depolarizing voltage step in each recording sweep. The current traces were filtered at 5 kHz, digitized at 10 kHz using a Digidata 1550b interface, and stored for off-line analysis. Next, recording was switched to current clamp. The resting membrane potential and the action potential were monitored for more than 5 minutes before drug applications. To induce action potentials, the neurons were commanded by multiple steps of hyperpolarization currents. Tetraethylammonium (TEA-Cl) and TTX (Sigma-Aldrich) were added to the artificial cerebrospinal fluid, to block K^+^ or Na^+^ channels, respectively. Electrophysiological recording data were first visualized with Clampfit 11 and exported to MATLAB (Mathworks, Natick, MA, http://www.mathworks.com) for analysis. The recording traces were visualized using Igor Pro 6.0 (WaveMetrics, Portland, OR, http://www.wavemetrics.com).

### Multi-Electrode Array (MEA)

After 4 days of differentiation, neurons were plated on poly-D-lysine/laminin coated CytoView MEA 24-well plates (Axion BioSystems, www.axionbiosystems.com) at a density of 5×10^4^ cells/well. Recordings from electrodes were made using a Maestro MEA system (Axion BioSystems). Data were sampled at 12.5 kHz, digitized, and analyzed using the Axion Integrated Studio software (Axion BioSystems) with a 200 Hz high pass and 4 kHz low pass filter and an adaptive spike detection threshold set at 6 times the standard deviation of the background noise for each electrode with 1 second binning. Total action potential counts, mean neuronal firing rates and total burst counts were quantified using the Axion Integrated Studio software.

For H_2_O_2_ and edaravone treatment, a 10-min MEA recording was acquired as the initial baseline at 0 h. Neurons were treated with edaravone followed by adding H_2_O_2_ to the edaravone-containing medium. Spontaneous action potential parameters after H_2_O_2_ treatment were normalized to the initial baseline.

### Neurotoxicity assay, Calcein-AM Staining, Neurite Tracing and High-content Analysis

Neurons (1 x 10^3^ per well) were plated in plates coated with poly-D-lysine/laminin with 10^3^ and 10^4^ per well for the 1,536- and 96-well plate, respectively. Neurons were pre-treated with edaravone (10 µM) for 16 hours in neurotrophic factor-free neurobasal medium followed by H_2_O_2_ (25 µM) or glutamate (200 µM) treatment for 24h. Live cell imaging used Calcein-AM dye (1 µM, Thermo Fisher Scientific). Neurite length quantification used the high-content analysis software (HCA-Vision V2.2.0. CSIRO). Neurite length per field was normalized to the number of Hoechst 33342 stained nuclei.

### RNA Sequencing and Quantitative Real-Time PCR (qPCR)

Total RNAs were extracted using the RNeasy Mini kit (Qiagen) and subjected to sequencing using the HiSeq 2500 platform (Illumina). Raw reads were aligned to reference human genome build hg19 using HISAT2 [22] with default parameters. For each gene, the number of reads aligned to its exons were counted and summarized into gene level counts by StringTie [23] based on the GENCODE hg19 annotation. Normalization between samples was carried out by the R package edgeR [24, 25], which controls sequencing depth and RNA composition effects. Heatmap was generated according to the count table with scaling across the samples for each gene. The RNA-Seq data sets can be accessed through the Gene Expression Omnibus (GEO) Repository (GSE151997).

qPCR analysis follow our previous publication [19]. Relative expression of each gene was normalized to the 18S rRNA. Primer sequences are listed in Table S6.

### Edaravone Treatment in **M**ice

C57BL/6J mice (Jackson Laboratory, 8 weeks old, female) received daily intraperitoneal administration of vehicle (saline) or edaravone (15 mg/kg body weight), following the previous publication [26]. Entire spinal cord tissues were harvested using the hydraulic extrusion method [27]. Total proteins were extracted using RIPA buffer containing protease and phosphatase inhibitors.

### Availability of Data and Materials

Further information and requests for resources and reagents will be fulfilled by the corresponding author. All unique/stable reagents generated in this study are available with a completed Materials Transfer Agreement. The RNA-Seq datasets reported here have been deposited to the Gene Expression Omnibus (GEO) Repository (GSE151997).

### Study Approval

All experiments involving human stem cells have been approved by the Johns Hopkins Medicine Institutional Review Board. The animal protocol was approved by the Johns Hopkins School of Medicine Animal Care and Use Committee.

### Statistics

All quantifications were performed by observers blinded to the experimental groups. All results represent at least three replicates with details in each figure legend. All data are represented as Mean ± SEM. Statistical analysis was performed using Prism software (GraphPad, San Diego, CA, http://www.graphpad.com). For comparing two groups, unpaired, two-tailed Student’s t test was performed (minimal requirement: *p* < 0.05). For more than two groups, one-way ANOVA with Tukey’s honestly significant difference (HSD) post-hoc test was used (minimal requirement: *p* < 0.05). qRT-PCR analysis used t test with the Bonferroni correction for multiple comparisons. Other statistical tests were specified in each figure legend.

## Results

### Synthetic mRNAs coding Olig2 and Ngn2 with phosphosite modification induce efficient MN differentiation from iPSCs

Ngn2 and Olig2 are two TFs co-expressing in motor neuron progenitor cells, and their ectopic expression in combination induces MNs in the chick neural tube [17]. Intrigued by these results, our goal is to develop synthetic mRNAs to ectopically express Ngn2 and Olig2 in iPSCs for efficient generation of mRNA-induced MNs, hereinafter referred to as miMNs. We previously reported that mRNAs coding Ngn2 with eight serine-to-alanine modifications (Figure 1A, referred to as N-SA) lead to higher protein expression and more efficient neuronal conversion in iPSCs, compared to mRNAs coding wild-type Ngn2 [19]. Thus, we use N-SA mRNAs here for MN differentiation. To optimize Olig2 mRNAs, we compared mRNAs coding wild-type Olig2 and a modified form with three serine-to-alanine mutations at S10, S13 and S14 sites (Figure 1A, referred to as O-WT and O-SA, respectively). We co-transfected iPSCs using N-SA mRNAs in combination with two forms of Olig2 mRNAs to determine Olig2 protein levels in this context with Ngn2 expression. O-SA mRNAs produced 2.6-fold and 3.7-fold more proteins than O-WT mRNAs, at 24h and 48h after transfection, respectively (Figure 1B), suggesting that O-SA mRNAs may more efficiently induce MN differentiation. We further tested six differentiation schemes (Figure 1C) that combine N-SA with O-WT or O-SA mRNAs, also including SHH, a known morphogen for MN lineage specification [28], and DAPT, a Notch signaling inhibitor widely used to promote neuronal conversion [29]. Three daily co-transfections of N-SA/O-SA mRNAs plus SHH/DAPT most efficiently induced MN lineage conversion, based on significantly higher levels of HB9 and Islet1, two well-defined MN lineage markers (Figure 1D). This combination is also among the top-three schemes showing equal efficiency in inducing the neuronal marker NeuroD1 (Figure 1D). In contrast, other schemes (e.g. N-SA/O-WT/SHH/DAPT) did not show similar efficiency in simultaneously inducing neuronal and MN lineage markers.

**Figure 1.**
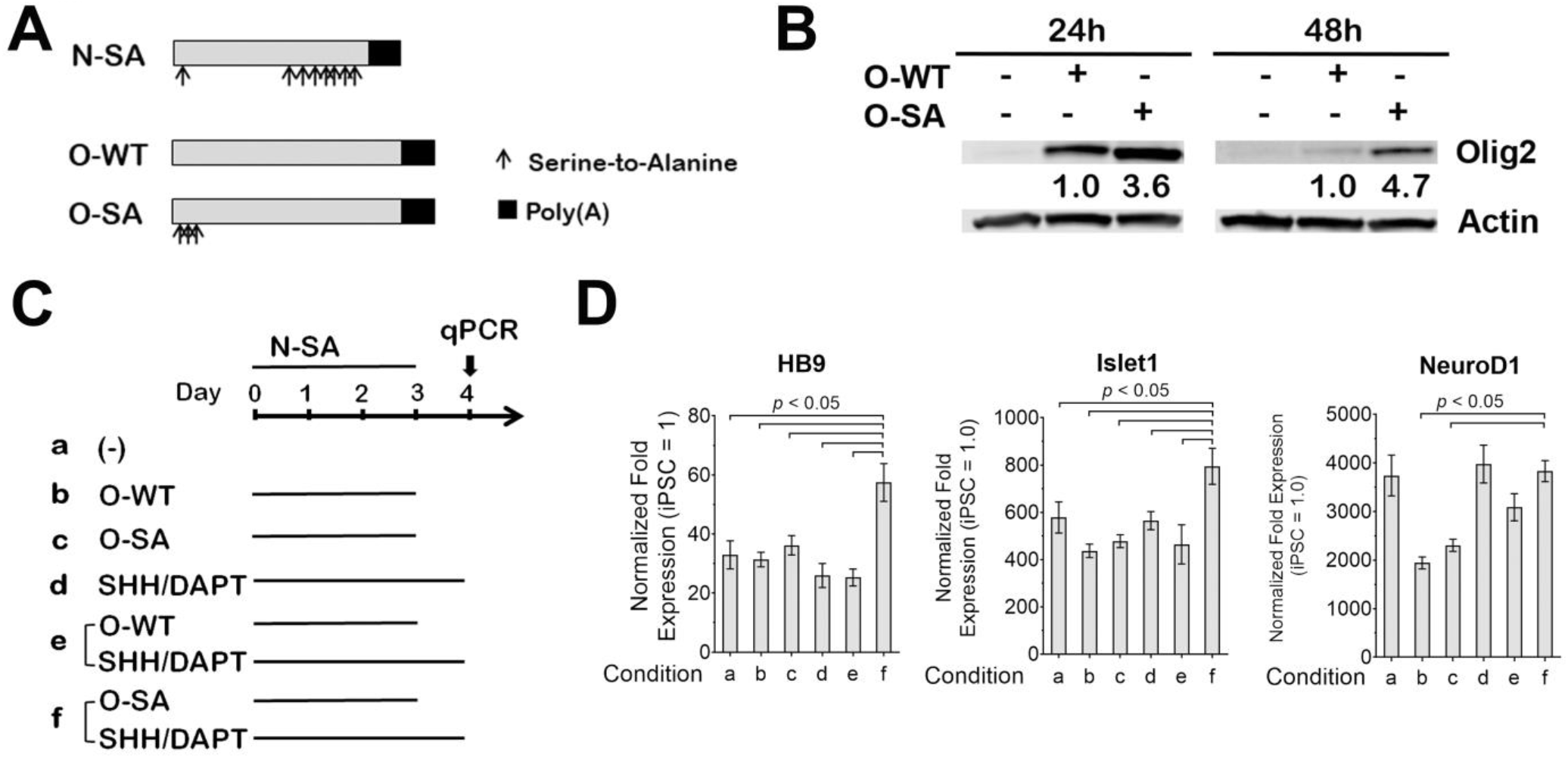
mRNAs coding phosphosite-modified Olig2 and Ngn2 induce efficient conversion of iPSCs to the MN lineage. **(A)** Diagram of mRNAs coding phosphosite-modified Ngn2 (N-SA), wild-type Olig2 (O-WT) and phosphosite-modified Olig2 (O-SA). Arrows indicate serine-to-alanine modifications (Ngn2: at 24, 193, 207, 209, 219, 232, 239 and 242 amino acids; Olig2: at 10, 13, 14 amino acids). **(B)** Normal iPSCs (iPSC1) received a single transfection of O-WT or O-SA mRNAs in combination with N-SA mRNAs. Total cellular proteins were harvested at 24 and 48h for Olig2 western blotting. Protein fold expression normalized to Actin are shown below each lane (O-WT samples = 1.0). **(C** and **D)** Schematic diagram of six conditions tested for identifying the most efficient strategy for MN lineage conversion. In all conditions, iPSC1 cells received three daily transfections of N-SA mRNAs in combination with O-WT, O-SA mRNAs and/or SHH/DAPT. Cells at day 4 of differentiation were subjected to qPCR to compare two MN lineage markers (HB9 and Islet1) and the pan-neuronal marker NeuroD1. Gene expression was normalized to the levels of undifferentiated iPSCs. Data represents Mean ± SEM from qPCR with 3 technical replicates, *p* values are calculated by one-way ANOVA with Tukey’s HSD pos-hoc test.

Guided by these results, we established a rapid 4-day MN differentiation protocol, including three daily co-transfections of N-SA/O-SA mRNAs plus SHH/DAPT treatment (Figure 2A, also see Materials and Methods). In 4 days, mRNA transfection converted iPSCs to TUJ1^+^ neuronal cells (Figure 2B, TUJ1^+^: >90%). These cells, after being passaged, can be cryopreserved or matured *in vitro* to miMNs showing typical neuronal morphology (Figure 2B, TUJ1^+^: >95%) and expressing various MN markers as characterized below. This protocol reproducibly generated miMNs from two normal iPSC lines (iPSC1 and iPSC3), an ALS iPSC line with the SOD1^A4V^ mutation (iPSC2), as well as human ESCs (Figure 2C).

**Figure 2.**
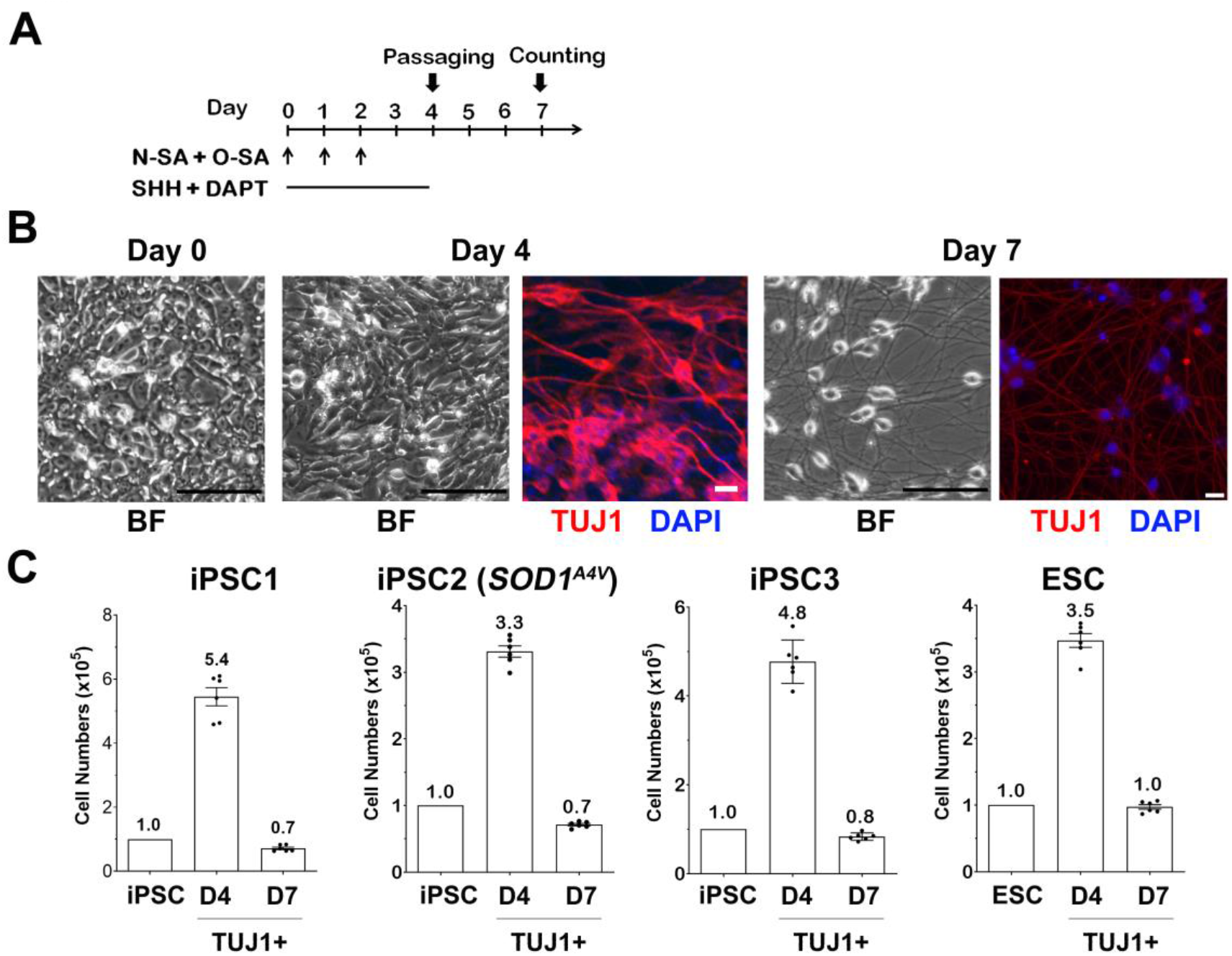
mRNA-induced MN generation from iPSCs. **(A)** Diagram of the differentiation protocol. **(B)** Brightfield (BF) microscopic images show iPSC1 cells and differentiated cells at indicated days (Bar: 100μm). Cells at day 4 and 7 of differentiation were immunostained for TUJ1 for quantification (Bar: 20μm; DAPI: nuclei). **(C)** iPSCs and ESCs (10^5^ cells) were differentiated by mRNAs. TUJ1^+^ cells at differentiation day 4 (before passaging) and 7 (after replating) were counted after TUJ1 immunostaining with cell numbers shown inside each panel. Data represents Mean ± SEM (n = 6, 3 independent differentiation, and each differentiation provides 2 wells of cells for immunostaining and quantification).

miMNs from normal and ALS iPSCs as well as ESCs express known MN markers, including HB9, Islet1 and the cholinergic neuron marker ChAT (Figure 3A and 3B; HB9^+^/TUJ1^+^: >92%; Islet1^+^/TUJ1^+^: >70%; ChAT^+^/TUJ1^+^: >97%). miMNs also co-express MN markers (Figure 3C; HB9^+^/ChAT^+^/ TUJ1^+^: >93%; Islet1^+^/ChAT^+^/TUJ1^+^: >65%). The pluripotent stem cell marker (OCT4) and the oligodendrocyte lineage marker (O4) were not detected in miMNs (Supplemental Figure 1A). The cholinergic neuron marker ChAT is more abundantly expressed by miMNs at day 20 of differentiation (Fig. 3A and 3B), compared to those at day 10 of differentiation (Supplemental Figure 1B). miMNs after *in-vitro* maturation also express the synaptic vesicle protein and mature neuron marker Synapsin 1 along TUJ1+ nerve fibers (Figure 3D). Ngn2 induction alone has been reported to induce non-MN subtypes in iPSCs and ESCs (e.g. cortical glutamatergic neurons) [30, 31]. Thus, we assessed markers for glutamatergic (VGluT1) and GABAergic (GAD67) neurons and showed the absence of these markers in miMNs (Figure 3E), supporting MN conversion driven by N-SA and O-SA in combination. As a control, iPSCs were differentiated by three daily transfections of NSA mRNA alone without using OSA mRNA or SHH/DAPT. NSA-induced TUJ1+ neurons express the glutamatergic marker VGlut1 (Supplemental Figure 1C), which result is consistent with publications showing glutamatergic neuron differentiation driven by Ngn2 alone [30, 31].

**Figure 3.**
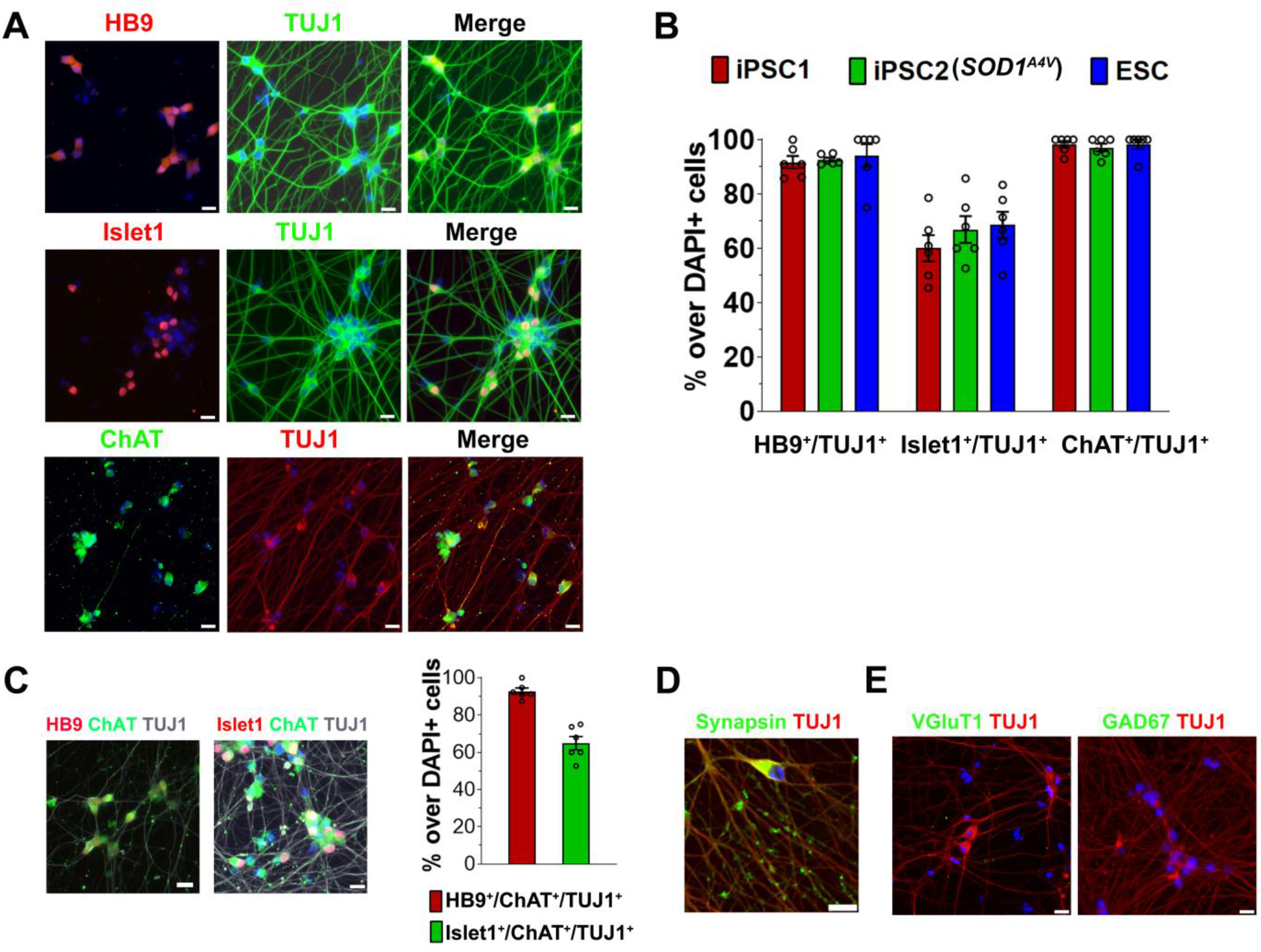
miMNs express pan-neuronal and MN lineage markers. **(A and B)** HB9^+^/TUJ1^+^, Islet1^+^/TUJ1^+^ and ChAT^+^/TUJ1^+^ neurons were detected by immunostaining and quantified in miMNs from the iPSC1, iPSC2 and ESC line. Neurons at day 10 (HB9 and Islet1) and 20 (ChAT) of differentiation were analyzed. **(C)** HB9^+^/ChAT^+^/TUJ1^+^ and Islet1^+^/ChAT^+^/TUJ1^+^ cells were quantified in miMNs from the iPSC1 line (day 20 of differentiation). **(D** and **E)** iPSC1-derived miMNs (day 30 of differentiation) were immunostained for the mature neuron marker Synapsin-1, glutamatergic marker VGlut1 and GABAergic marker GAD67. Cell nuclei were counterstained with DAPI (Bar = 20 μm). Data represents Mean ± SEM (n = 6, 3 independent differentiation, and each differentiation provides 2 wells of cells for immunostaining and quantification).

### Electrophysiological and high-content analysis of miMNs

The functional maturation of miMNs was studied using patch-clamp recording (Figure 4A). In miMNs from the normal iPSC line iPSC1, all of 36 recorded neurons (15-20 days of in-vitro maturation) showed spiking following steps of hyperpolarization current injections (Figure 4B). Repetitive multiple action potentials were induced by the hyperpolarization in 19 of 36 neurons (Figure 4C, left panel). The action potentials were confirmed by blocking with TTX, a selective sodium channel blocker [32] (Figure 4C). As generation of action potentials requires a coordinated interplay between sodium and potassium channels, we showed that TEA, a voltage-dependent K^+^ channel blocker [33] attenuated the repetitive action potentials that recovered after compound washing-out (Figure 4D). In addition, the late interspike intervals (ISIs) were significantly longer than that for the preceding ISIs (Figure 4E; 24.3 ± 1.3 vs 13.8 ± 1.2 ms, *p* < 0.001), implicating that miMNs exhibit adaptation to extending hyperpolarization, a previously reported physiological property of *bona fide* MNs [34–36].

**Figure 4.**
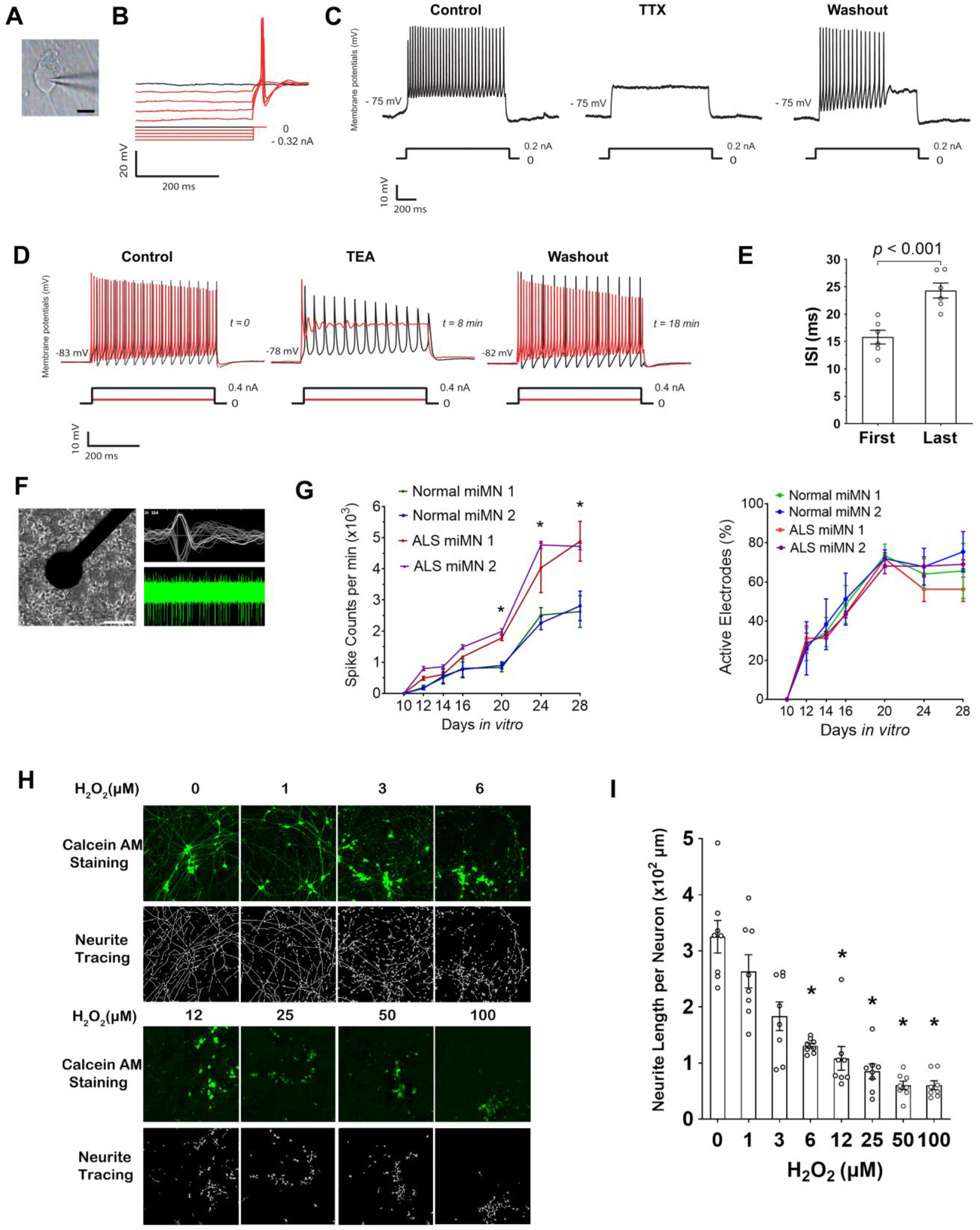
Functional characterization and high-content analysis of miMNs. **(A)** Microscopic image of a patched miMN (Bar = 10 μm). **(B)** Typical rebound potentials after steps of hyperpolarization in miMNs. **(C** and **D)** miMNs displayed repetitive action potentials after depolarization commands. The action potentials were reversibly blocked by TTX (C) and TEA (D). Traces underneath each panel indicate current steps. **(E)** Interspike intervals (ISI) to depolarization current injections in miMNs are significantly delayed (n = 6, *p* = 0.001, t-test). The first ISI indicates the ISI between the first and second spike following depolarization commands, whereas the last ISI indicates the ISI between the last two spikes after depolarization commands being withdrawn. **(F)** A brightfield image shows miMNs attached on the electrode underneath the MEA plate (Bar = 100 µm). The right panels show spiking waveforms (top) and spontaneous spiking activity of miMNs (bottom) recorded by one electrode. **(G)** Normal miMNs from iPSC1 and iPSC3 iPSCs (referred to as normal miMN 1 and 2, respectively) and ALS miMNs from iPSC2 and iPSC4 iPSCs (referred to as ALS miMN 1 and 2, respectively), were plated to the MEA plate at day 4 of differentiation. Their spontaneous spiking was recorded at indicated days *in vitro*. ALS miMNs showed more spiking activity than normal miMNs. (3 technical replicates for each miMN line, *: *p* < 0.05, linear regression with clustered data, ALS vs normal miMNs) **(H** and **I)** miMNs from iPSC1 at day 4 of differentiation were cultured in the 1,536-well plate for 2 days, and subjected to 24h H_2_O_2_ treatment followed by Calcein AM staining, neurite tracing (H) and neurite length quantification (I). Neurite length was normalized to nuclei numbers (6 wells for each condition as technical replicates, *: *p* < 0.01, one-way ANOVA with Tukey’s HSD pos-hoc test, compared to the untreated control) Data represents Mean ± SEM.

Next, we applied the microelectrode array (MEA) system to continuously monitor spontaneous neuronal activities of miMNs in a high-throughput manner. miMNs formed direct contact with electrodes for recording spontaneous spikes (Figure 4F). During *in-vitro* maturation, normal miMNs from two iPSC lines showed more active spontaneous firing and their spiking rate reached the peak after 24 days (Figure 4G, left). ALS miMNs from two iPSC lines with the *SOD1^A4V^* mutation showed more active spontaneous firing than normal miMNs (Figure 4G, left, *p* < 0.05), consistent with the previous report showing hyperexcitability of MNs from ALS iPSCs [37]. The number of active electrodes did not show difference between normal and ALS miMNs (Figure 4G, right). At the end of the MEA analysis (30 days *in vitro*), neuron numbers in MEA wells showed no difference between normal and ALS miMNs (0.8-1.3 x 10^4^ per well).

miMNs are applicable to high-content analysis in the 1536-well format. miMNs from iPSC1 at day 4 of differentiation were plated using a microplate dispenser. After 48h, miMNs stained with Calcein-AM showed robust neurite outgrowth (Figure 4H). These miMNs were treated with hydrogen peroxide (H_2_O_2_) that has been used to model oxidative stress and neurotoxicity in ALS and other neurodegenerative diseases [38]. H_2_O_2_ dose-dependently induced morphological signs of neurotoxicity, including neurite fragmentation and cell body condensation (Figure 4H and 4I, IC_50_ = 5.4 µM, *p* < 0.01, compared to the untreated control).

### Edaravone protects miMNs from H_2_O_2_-induced neurotoxicity

We applied miMNs to explore the cellular and molecular effects of edaravone, an FDA-approved drug for ALS. In H_2_O_2_-induced neurotoxicity assay, edaravone significantly alleviated neurite damage in miMNs (Figure 5A and 5B). Edaravone-treated miMNs showed only 26% reduction of neurite length after H_2_O_2_ (25 µM) treatment, compared to the edaravone-untreated control (*p* < 0.05). This neurite damage was less severe than the 93% reduction of neurite length shown in edaravone-untreated neurons (*p* < 0.01). Without H_2_O_2_ treatment, edaravone did not significantly alter neurite length in miMNs. The neuroprotective effect of edaravone was further tested using the MEA system. We used H_2_O_2_ at a low concentration (3 µM) that does not significantly damage neurites as shown in Figure 4H and 4I). This low-dose H_2_O_2_ treatment still inhibited spontaneous spiking of miMNs by about 70% (Figure 5C and 5D), recapitulating the early impairment of neuronal functions before neurodegeneration. In H_2_O_2_-treated miMNs, edaravone effectively restored neuronal spiking to the same level as H_2_O_2_-untreated neurons (Figure 5C and 5D, *p* < 0.05). In H_2_O_2_-untreated miMNs, edaravone did not significantly alter spontaneous spiking, indicating that this drug may function through neuroprotection but not through elevating neuronal activity.

**Figure 5.**
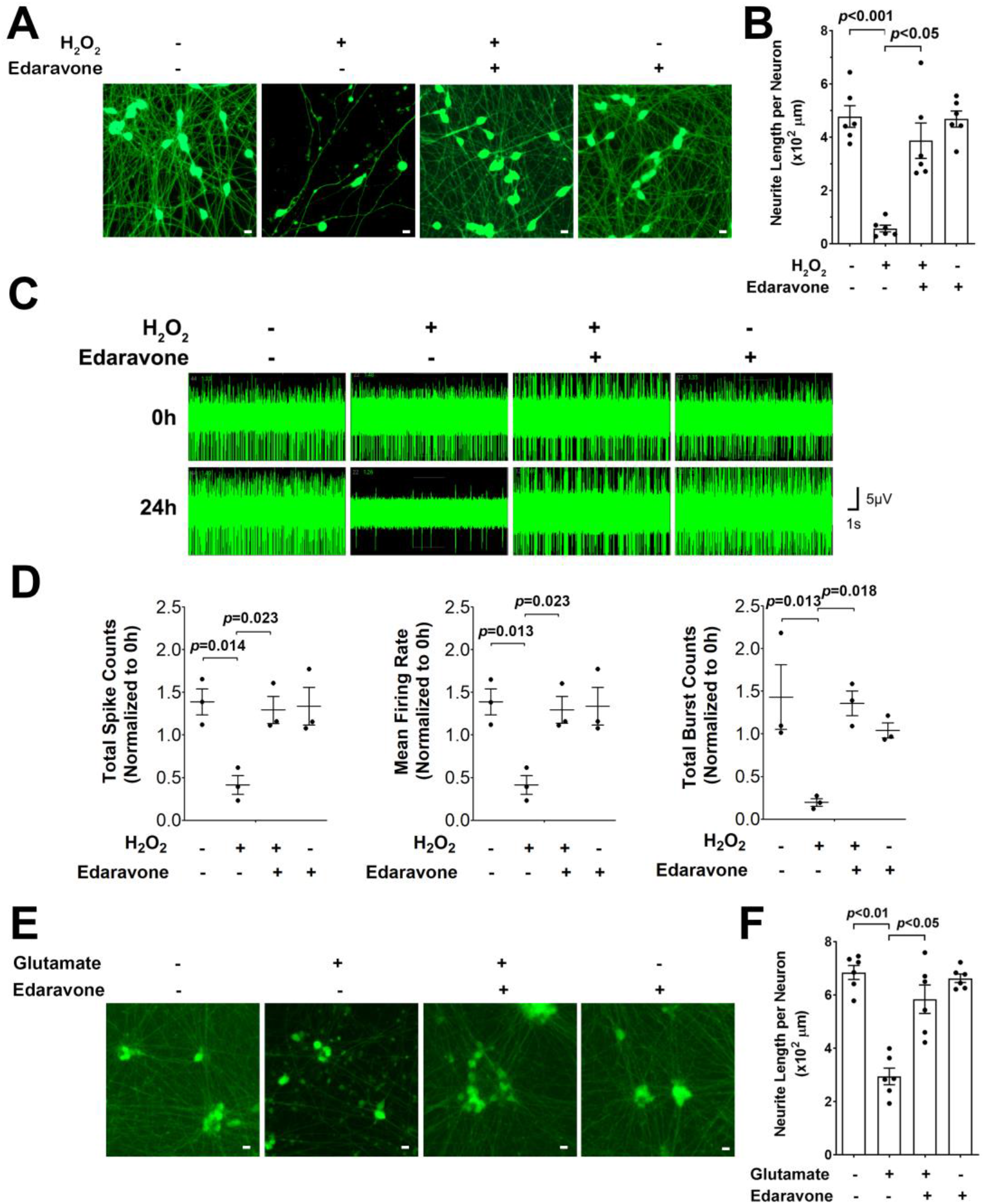
Edaravone protects miMNs from H_2_O_2_-induced neurotoxicity. **(A and B)** iPSC1-derived miMNs (day 7 of differentiation) were pre-treated with edaravone (10 µM) for 16 hours in neurotrophic factor-free medium followed by H_2_O_2_ treatment (25 µM, 24h). After Calcium AM staining (A, Bar = 10 µm), neurite length was quantified (B, 6 wells for each condition as technical replicates). Edaravone rescues H_2_O_2_-induced neurite damage. **(C** and **D)** iPSC1-derived miMNs (day 25 of differentiation) were pre-treated with edaravone (10 µM) in neurotrophic factor-free medium for 16 hours followed by H_2_O_2_ treatment (3 µM, 24h). Representative traces (C) show spontaneous spiking activity further quantified in D (3 wells for each condition as technical replicates). Edaravone restores spontaneous spiking activity impaired by H_2_O_2_. (**E** and **F**) iPSC1-derived miMNs (day 7 of differentiation) were pre-treated with edaravone (10 µM) for 16 hours in neurotrophic factor-free medium followed by glutamate treatment (200µM, 24h). After Calcium AM staining (E, Bar = 10 µm), neurite length was quantified (F, 6 wells for each condition as technical replicates). Edaravone rescues glutamate-induced neurite damage. Data represents Mean ± SEM with *p* values (one-way ANOVA with Tukey’s HSD pos-hoc test) shown inside each panel.

We also studied glutamate-induced neurotoxicity in miMNs and found that edaravone significantly alleviated glutamate-induced neurite damage in miMNs (Figure 5E and 5F). Edaravone-treated miMNs showed only 15% reduction of neurite length after glutamate treatment (200 µM, 24h), significantly less than the 57% reduction in edaravone untreated neurons (*p* < 0.05). Taken together, these results support the neuroprotective function of edaravone in miMNs, thus warranting studies to reveal underlying mechanisms.

### Transcriptomic profiling of edaravone-induced molecular responses in miMNs

We performed RNA-sequencing to identify differentially-expressed (DE) genes and their associated signaling pathways. The normal iPSC line (iPSC1) was independently differentiated into two sets of miMNs as technical replicates. After *in-vitro* maturation for 20 days, these two sets of miMNs were treated with edaravone (10 μM) or DMSO as the control for 24h. This edaravone concentration has been shown to protect miMNs from H_2_O_2_- and glutamate-induced neurotoxicity (Figure 5). Over 20 million cDNA reads were generated for each of the two conditions (two technical replicates for each condition) and showed a more than 90% alignment rate to the human genome. Consistency between two replicates was demonstrated by heatmap clustering (Supplemental Figure 2A). We also performed transcriptomic comparison between miMNs and MNs differentiated by the traditional compound-based method [39]. miMN samples more closely cluster with compound-induced MNs, especially those at 8-9 days of differentiation (Supplemental Figure 2B), compared to cells at the undifferentiated (Day 0) and early-induction (Day 1-2) stage.

There are 2,329 up-regulated (Table S1) and 1,916 down-regulated genes (Table S2) altered by edaravone (Figure 6A, FDR ≤ 0.01, log_2_(Fold-Change) Cut-off = ±0.5, see Table S3 for expression data of all detected genes). The Ingenuity Pathway Analysis (IPA) was used for gene functional annotation and pathway enrichment assay. IPA pathways enriched in up- and down-regulated genes were ranked in Table S4 and S5, respectively. Top ten IPA pathways (Figure 6B) include those associated with neuron functions and ALS pathogenesis, such as the synaptogenesis and CREB signaling enriched in up-regulated genes, and the mitochondrial dysfunction and oxidative phosphorylation signaling enriched in down-regulated genes. Four IPA pathways and associated DE genes were chosen for further studies (Figure 6C with highlighted genes for validation), including the ALS, superoxide radical degradation, GDNF and neurotrophin/TRK signaling pathways. This selection is based on essential roles of these pathways and their associated genes in MN survival and function, and ALS pathogenesis.

**Figure 6.**
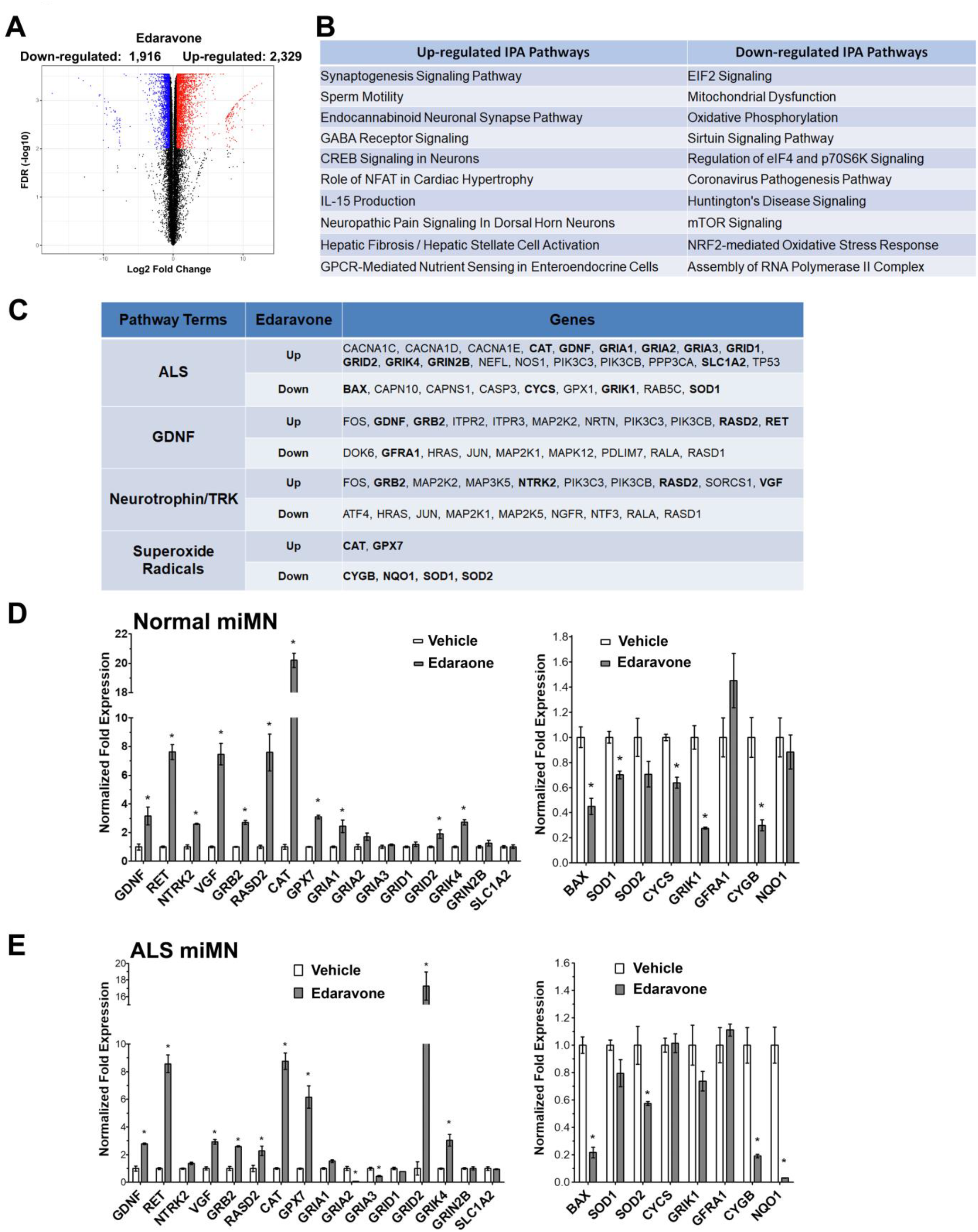
Transcriptomic analysis of edaravone-induced responses in miMNs. **(A)** Volcano plot showing DE genes after edaravone treatment in normal miMNs from iPSC1 (red and blue dots: genes with FDR ≤ 0.01 and log_2_(fold change) ≥ 0.5 or ≤ -0.5, respectively). **(B)** Top ten IPA pathways enriched in up- and down-regulated genes. Pathways are ranked based on - log(*p* value) as calculated by the Fisher’s exact test. **(C)** IPA pathways of interest and their associated genes. Genes selected for qPCR validation are highlighted. **(D** and **E)** Normal and ALS miMNs were treated with edaravone (10 µM, 24h) or DMSO as the vehicle control. Total cellular RNAs were subjected to qPCR analysis of up-regulated (left panel) and down-regulated (right panel) genes highlighted in C. Data represents Mean ± SEM from 3 technical replicates (*: *p* < 0.05, t test with the Bonferroni correction for multiple comparisons).

We used qPCR to validate DE genes of interest in both normal and ALS miMNs (Figure 6D and 6E). Validated up-regulated genes include key components of the neurotrophic factor signaling, such as *GDNF* and its receptor *RET*, another neurotrophic gene *VGF*, as well as two downstream signaling molecules (*GRB2* and *RASD2*). The GDNF co-receptor *GFRA1* is a down-regulated gene in the RNA-seq result, whereas qPCR did not detect its change in two miMN models. Validated up-regulated genes also include two antioxidant enzymes (*CAT* and *GPX7*). The up-regulation of some ALS signaling-associated genes was validated in normal and ALS miMNs, including *GRID2* and *GRIK4* from the glutamate receptor family. *GRIA1* upregulation is validated in normal miMNs. Down-regulated genes that were consistently validated in normal and ALS miMNs include the pro-apoptotic gene *BAX* and *CYGB* coding a stress-responsive hemoprotein expressed in the brain [40]. *SOD1* showed a small (15%) but significantly down-regulation in normal but not ALS miMNs. *SOD2* showed about 40% down-regulation only in ALS miMNs.

### Edaravone promotes the GDNF/RET neurotrophic signaling pathway in miMNs and the mouse spinal cord

Consistent with qPCR results, edaravone elevated the levels of RET and VGF proteins from the neurotrophic factor signaling and two antioxidant enzymes (CAT and GPX7), in normal and ALS miMNs (Figure 7A). Protein levels of the GDNF co-receptor GFRA1 were also elevated by edaravone in both miMN models (Figure 7A), although GFRA1 mRNAs were not altered by edaravone as previously determined by qPCR (Figure 6D and 6E). Next, we focused on the previously unrecognized effect of edaravone on the GDNF/RET neurotrophic signaling. Edaravone treatment in normal and ALS miMNs induces the expression of HOXB5, NKX2.1 and PHOX2B (Figure 7B), three TFs transactivating the RET promoter [41, 42], suggesting the involvement of these TFs in activating RET transcription by edaravone. As a control, we tested other TFs (CTCF, MYC, RAD21 and GABPA), the binding motifs of which are found in the RET promoter. We did not detect their induction by edaravone (data not shown). To determine the functional outcome of GDNF receptor induction by edaravone, we performed GDNF stimulation (30 min) in normal and ALS miMNs that have been cultured overnight in neurotrophic factor-free medium with or without edaravone. Edaravone-treated miMNs showed higher levels of total and phosphorylated RET proteins than the untreated control (Figure 7C). More importantly, edaravone-treated miMNs showed higher levels of phosphorylated AKT, ERK and Src (Figure 7C), three GDNF/RET-activated downstream kinases [43]. These results support that edaravone-induced RET expression leads to more active GDNF/RET signaling. We further asked if edaravone-induced RET expression leads to enhanced neuroprotection. We used a high dose of H_2_O_2_ treatment (50 µM, 6h) to induce severe neurite damage in normal miMNs (Figure 7D, about 95% neurite loss), which cannot be effectively protected by edaravone or GDNF alone (Figure 7D, *p* >0.05). In comparison, edaravone/GDNF combined treatment significantly inhibited H_2_O_2_-induced neurite damage (Figure 7D, neurite loss: 37% compared to 95%, *p* < 0.01). Consistently, edaravone/GDNF combined treatment also protected ALS miMNs with the *SOD1^A4V^* mutation (Figure 7E, neurite loss: 18% compared to 91%, *p* < 0.01), more effectively than their single treatment (neurite loss: 58% and 72% for GDNF and edaravone single treatment, respectively).

**Figure 7.**
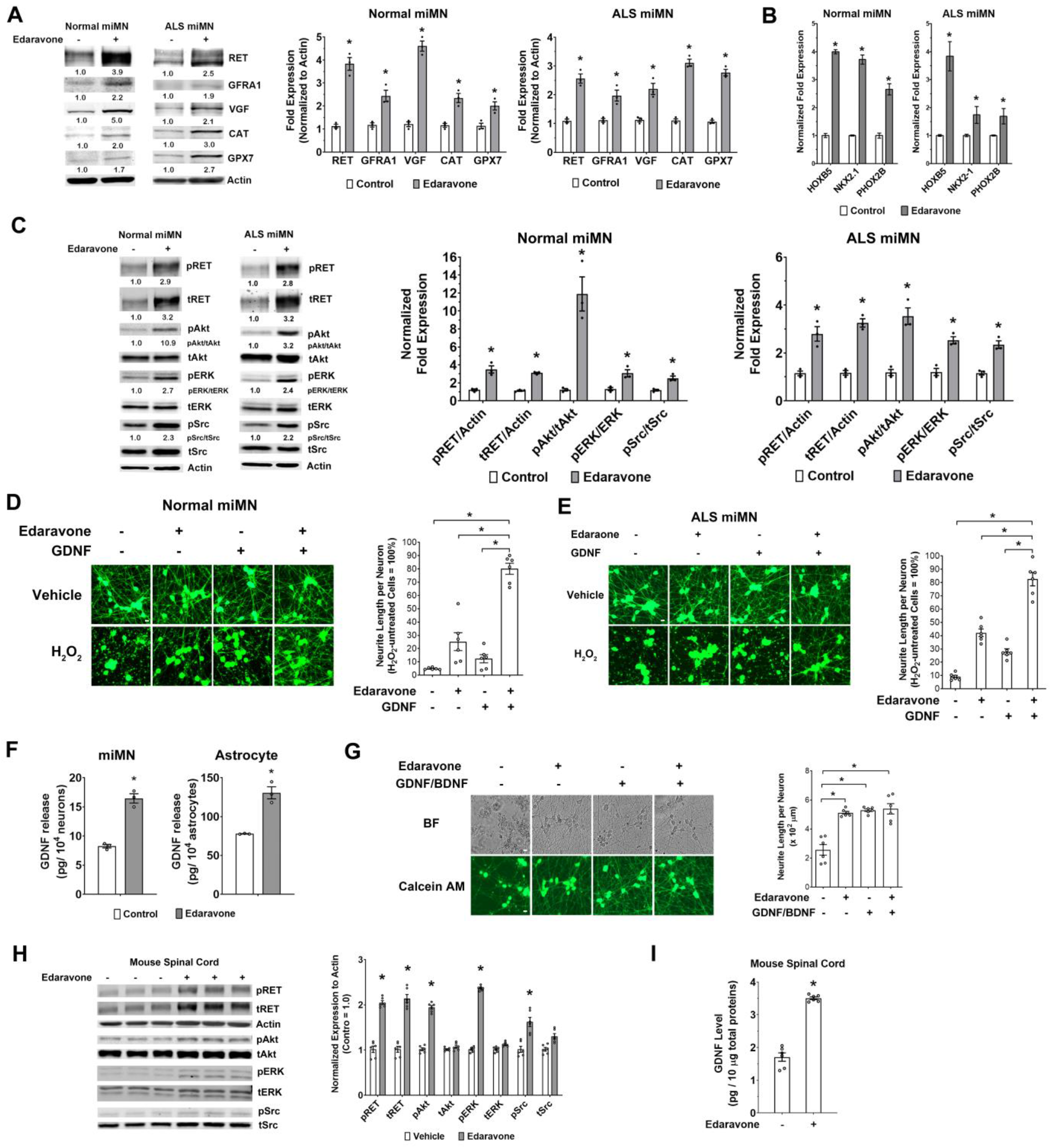
Edaravone induces the GDNF/RET neurotrophic signaling pathway in miMNs and the mouse spinal cord. **(A and B)** Normal and ALS miMNs were subjected to edaravone treatment (10 µM, 24h) or DMSO as the vehicle control. Total cellular proteins were subjected to western blotting of genes for validation (A, left) and quantification from three blotting results (B, right). Total cellular RNAs were subjected to qPCR analysis of three TF genes (B). (*: *p* < 0.05, t test with bonferroni correction for multiple comparisons) **(C)** Normal and ALS miMNs with +/- edaravone treatment (10 µM, 24h) were subjected to western blotting (left) and quantification from three blotting results (right) to measure the levels of GDNF/RET signaling components (*: *p* < 0.05, t test with the Bonferroni correction for multiple comparisons). **(D** and **E)** Normal and ALS miMNs (day 7 of differentiation) were pre-treated with +/- edaravone (10 µM) and GDNF (1 ng/ml) for 16h in neurotrophic factor-free medium and were treated with H_2_O_2_ (50 µM) or PBS as the vehicle control for 24h. Neurite length was quantified from 6 wells for each condition as technical replicates, after Calcium AM staining (Bar = 10 µm). Edaravone+GDNF more effectively alleviated H_2_O_2_-induced neurite damage than single treatment. **(F)** Normal miMNs and astrocytes differentiated from human NSCs were subjected to edaravone treatment (10 µM) for 48h. GDNF release in the culture medium was measured by ELISA with 3 technical replicates. **(G)** Normal miMNs were cultured in neuron maturation medium with +/- edaravone and GDNF/BDNF for 15 days. Brightfield (BF) and Calcium AM staining images were shown (Bar = 10 µm). Neurite length from 6 wells for each condition as technical replicates was compared among different conditions. **(H** and **I)** Mice (n=6 for each group) received intraperitoneal injection of edaravone (15 mg/kg daily) or vehicle (saline) for 5 days. Total proteins from spinal cord tissues were analyzed by western blotting (G) Representative western blotting and quantification of protein fold expression normalized to Actin (*: *p* < 0.05, t test with the Bonferroni correction for multiple comparisons). Total proteins from spinal cord tissues from 6 mice were also analyzed by GDNF ELISA (H, *: *p* < 0.01, t test). Data represents Mean ± SEM.

RNA-seq and qPCR results both show that GDNF transcription is also induced by edaravone (Figure 6). Consistently, we found that edaravone increased GDNF release from miMNs by about 100% (Figure 7F, *p* < 0.05). In astrocytes differentiated from human fetal neural stem cells (NSCs), edaravone also increased GDNF release by 68% (Figure 7F, *p* < 0.05). These results indicate that edaravone-induced GDNF release may lead to more active GDNF/RET neurotrophic signaling in autocrine and possibly also paracrine manners.

Since edaravone-induced RET expression and GDNF release may provide autocrine neurotrophic signaling to support neuron survival and maturation, we tested if edaravone can substitute the neurotrophic factors (GDNF and BDNF) used in miMN maturation medium. After 15 days of *in-vitro* maturation, we found that medium containing edaravone alone, GDNF/BDNF or both showed no difference in supporting normal neuronal morphology and neurite outgrowth (Figure 7G). miMNs from these three conditions also showed no difference in the expression of mature MN markers (e.g. ChAT and Synapsin 1) (data not shown). In contrast, miMNs survived poorly in medium without GDNF/BDNF or edaravone, and showed impaired neurite outgrowth (Figure G). These results further support the neurotrophic function of edaravone-induced GDNF/RET signaling in miMNs.

Finally, we tested the *in-vivo* effect of edaravone on the GDNF/RET signaling in the spinal cord of mice with systemic edaravone treatment. Mice received intraperitoneal edaravone or vehicle treatment for 5 days. By analyzing total proteins from the entire spinal cord, we showed that edaravone treatment, compared to vehicle control, increased the levels of total and phosphorylated RET proteins by about 100%, and led to enhanced activation of RET downstream kinases (AKT, ERK and Src) (Figure 7I). GDNF levels in mouse spinal cord tissues also showed more than 100% increase after edaravone treatment, as measured by ELISA (Figure 7J). Overall, these results support edaravone’s *in-vivo* efficacy to activate the GDNF/RET neurotrophic signaling.

## Discussion

Synthetic mRNAs are highly efficient vehicles for delivering TF drivers to manipulate cell identity. Currently, mRNA-driven cell reprogramming kits are widely used for generating research and clinical grade human iPSCs [44]. Compared to cell reprogramming, mRNA-driven cell differentiation requires more optimization, including TF driver selection, TF sequence modification and mRNA delivery optimization, because a practical iPSC differentiation strategy requires at least 80% efficiency, compared to the 1-2% efficiency of cell reprogramming that is usually adequate [45, 46].

Traditional MN differentiation strategies for iPSCs use multiple compounds in a multi-step protocol and take 10-14 days to generate MNs with variable purity (commonly 50%-70%) [15, 16]. Here, we established a 4-day one-step protocol to reproducibly generate miMNs with >90% purity from normal and ALS patient-derived iPSC lines as well as ESCs. We defined Ngn2 and Olig2 as a pair of TFs sufficient to drive highly efficient MN conversion. It is noteworthy to mention that Olig2 alone is not capable of inducing neuronal conversion. Compared to the Ngn2/Olig2 combination, a previous report used mRNAs coding five TFs including wild-type Ngn2 to generate human iPSC-derived MNs in 7 days [47]. Mazzoni et. al. used three TFs (Ngn2, Isl1 and Lhx3) to differentiate mouse ESCs to MNs [18]. In the context of mRNA-based TF delivery, our results support that phosphosite modification in Ngn2 and Olig2 is a key element to increase the efficiency of mRNA-induced TF expression and MN conversion, as phosphosite modification leads to higher and long-lasting expression of Ngn2 and Olig2. SHH and DAPT were also found to promote MN induction, as MN lineage markers (HB9 and Islet1) were further induced by SHH/DAPT when N-SA/O-SA mRNAs were used. It is possible that mRNAs coding other TFs can be added to this protocol, which optimization may promote *in-vitro* survival and functional maturation of miMNs. Now, large-scale mRNA synthesis is easy and cost-effective. It is also noteworthy to mention that mRNAs used here were synthesized using nucleotides without pseudouridine and 5-methylcytidine modification, further reducing the cost of this method. In addition, miMNs after cryopreservation have a typical recovery rate of >60%. Therefore, it is practical to establish a bank of iPSC-derived miMNs from familial and sporadic ALS patients, which platform will facilitate drug development for this devastating disease with complex biology and significant clinical heterogeneity [4]. We extensively characterized the identity of miMNs using well-defined MN markers as well as transcriptomic comparison with MNs differentiated by the traditional compound-based protocol [39]. Furthermore, we characterized their neuronal functions, using traditional patch-clamping recording as well as the MEA system suitable for high-throughput analysis. Moreover, high-content analysis of miMNs in the 1536-well format will allow ultra-high-throughput drug screening in an ALS MN bank, especially drug combination screening. It is noteworthy to mention that transcriptomic similarity between miMNs and compound-induced MNs at different maturation stages may be altered by batch effects in RNA-seq data. Thus, expression of mature neuron markers (e.g. Synapsin-1) and electrophysiological characterization may more reliably define the maturation status of miMNs.

Our following study of edaravone supports the applicability of miMNs in ALS drug development. Edaravone effectively protects miMNs from oxidative stress-associated neurotoxicity, consistent with its clinical efficacy shown in some ALS patients and proposed drug action as a free-radical scavenger to reduce oxidative stress. We also showed the neuroprotective function of edaravone in gluatamate-induced neurotoxicity model using miMNs. Notably, the MEA analysis provides a more sensitive readout of neuronal damage as reflected by loss of neuronal activity. Since H_2_O_2_ is used at a much lower concentration without inducing rapid neurite damage, this MEA assay may more closely recapitulate a physiologically relevant condition of oxidative stress and neuronal dysfunction commonly seen during early ALS pathogenesis [4]. Compared to traditional screening platforms using neuron death or neurite damage as the readout, MEA analysis will help us identify potential compounds protecting neuronal functions of MNs and likely other neuron types involved in neurodegenerative disorders. To our best knowledge, this study provided the first genome-wide map of edaravone-regulated signaling pathways in human MNs from iPSCs. This transcriptomic study will provide the foundation to explore mechanisms of action of edaravone and facilitate its clinical applications in various brain disorders. A significant discovery is edaravone-activated neurotrophic factor signaling pathways, especially GDNF/RET signaling. Edaravone induces the expression of GDNF receptors (e.g. RET) as well as GDNF release in miMNs, supporting by following results showing more active RET downstream kinases in edaravone-treated miMNs. Edaravone also induces GRB2 and RASD2, two key signaling molecules of the GDNF/RET signaling [43], which mechanism may further boost the activation of this neurotrophic signaling. GDNF is a potent neurotrophic factor promoting MN survival, and dysfunction of GDNF/RET signaling plays an essential role in ALS pathogenesis [48]. GDNF-based ALS therapies have been extensively studied in pre-clinical ALS models [49, 50], including recombinant GDNF proteins, GDNF gene therapy and GDNF-secreting cells. Some of these therapies have been advanced to human trials, such as GDNF-secreting neural progenitor cells (NCT02943850), but still need optimization. Our results demonstrate that the brain-penetrating ALS drug edaravone promotes RET expression and GDNF release in MNs, which mechanism likely mediates the neuroprotective effect of edaravone on miMNs. RET down-regulation has been found in MNs of the *SOD1^G93A^* ALS mouse model. It is possible that ALS therapies simply supplying GDNF may be less effective than a therapeutic strategy that boosts both RET expression and GDNF supply. To support this hypothesis, we showed that edaravone and GDNF combined treatment more effective protect miMNs than their single treatment, which result justifies further pre-clinical and clinical studies of this combination strategy. Strikingly, edaravone can substitute all neurotrophic factors (GDNF/BDNF) in the miMN maturation medium to support *in-vitro* survival and maturation of miMNs, providing another evidence to support its neurotrophic function. To our best knowledge, this is the first report defining a small-molecule compound capable to replace neurotrophic factors essential for neuron culture. We also show that edaravone induces GDNF release in astrocytes, a major GDNF-secreting cell type in the brain, suggesting an edaravone-activated paracrine mechanism of neuroprotection. Moreover, edaravone shows consistent *in-vivo* efficacy to activate the GDNF/RET signaling in mouse spinal cord samples, justifying more studies in mouse models of ALS and other neurodegenerative diseases that may benefit from neuroprotection through GDNF/RET activation. Overall, our results provide a pharmacological strategy to activate the GDNF/RET neuroprotective signaling using a brain-penetrating drug with validated oral bioavailability.

The edaravone-induced neurotrophic protein VGF is a potential biomarker of ALS progression. VGF levels in the cerebral spinal fluid (CSF) and serum progressively decline in the *SOD1^G93A^* ALS mouse model and ALS patients, and exogeneous VGF is neuroprotective in MN cultures and ALS mouse models [51–53]. VGF is known to be induced by neurotrophic factors (e.g. BDNF and NGF) [54, 55] and also shows up-regulation after RET activation in neuroblastoma cells [56]. Thus, it is possible that edaravone induces VGF through the activation of GDNF/RET and/or BDNF/NTRK2 signaling. Elevated VGF levels in edaravone-treated miMNs may also function as a neuroprotective mechanism. Importantly, it is feasible to measure VGF-derived peptides in CSF and blood samples [51, 57]. Edaravone-induced VGF may be developed as a biomarker to predict drug response in clinical trials, such as the current ALS trial studying oral edaravone (NCT04165824).

Other edaravone-regulated genes with potential ALS association are also of interest, such as enzymes clearing superoxide radicals (e.g. CAT and GPX7), glutamate receptors (e.g. GRIA1, GRID2 and GRIK4) and pro-apoptotic proteins (e.g. BAX). More studies are required to understand how edaravone modulates various glutamate receptors that either positively or negatively regulate glutamate receptor-mediated excitotoxicity, a key pathogenetic mechanism in ALS [4, 58]. It is noteworthy to mention that our MEA analysis did not detect hyperexcitability in edaravone-treated miMNs.

## Conclusion

We established a synthetic mRNA-driven strategy to efficiently generate iPSC-derived functional MNs applicable to high-throughput drug screening. We further define the neuroprotective effect of the ALS drug edaravone and reveal underlying molecular mechanisms, including the activation of GDNF/RET neurotrophic signaling. Methodology from this study will expand the applications of iPSC technology in MN research and therapeutic development for MN-associated neurodegenerative diseases. Novel molecular insights from this study will facilitate the development of edaravone-based therapies (e.g. edaravone/GDNF combination therapy) for ALS and likely other brain disorders associated with neurodegeneration (e.g. Parkinson’s disease and stroke).

## List of abbreviations

iPSC: induced pluripotent stem cell
ESC: embryonic stem cell
PSC: pluripotent stem cell
MN: motor neuron
ALS: amyotrophic lateral sclerosis
miMN: mRNA-induced motor neuron
FDA: the Food and Drug Administration
TF: transcription factor

## Declarations

### Consent for publication

Not applicable.

### Availability of supporting data

The RNA-Seq datasets reported here have been deposited to the Gene Expression Omnibus (GEO) Repository (GSE151997).

## Authors’ contributions

All authors read and approved the final manuscript.

Q.L.: collection of data, data analysis and interpretation, manuscript writing, final approval of manuscript.

Y.F.: collection of data, data analysis and interpretation, final approval of manuscript.

Y.X.: collection of data, data analysis and interpretation, manuscript writing, final approval of manuscript.

X.Z.: collection of data, data analysis and interpretation, manuscript writing, final approval of manuscript.

Y.F.: collection of data, data analysis and interpretation, final approval of manuscript.

G.G.: data analysis and interpretation, manuscript writing, final approval of manuscript.

W.Z.: data analysis and interpretation, manuscript writing, final approval of manuscript.

J.R.: provision of study material, final approval of manuscript.

A.T.: provision of study material, final approval of manuscript.

P.L.: provision of study material, final approval of manuscript.

X.M.: provision of study material, final approval of manuscript.

N.M.: conception and design, provision of study material, manuscript writing, final approval of manuscript.

M.Y.: conception and design, financial support, collection of data, data analysis and interpretation, manuscript writing, final approval of manuscript.

## Acknowledgements

We thank Dr. Ted Dawson and Dr. Valina Dawson for helping our studies of human pluripotent stem cells. We thank Dr. John Laterra for helpful discussion and editing the manuscript.

## Authors’ information

None.

---

**Supplemental Figure 1.**
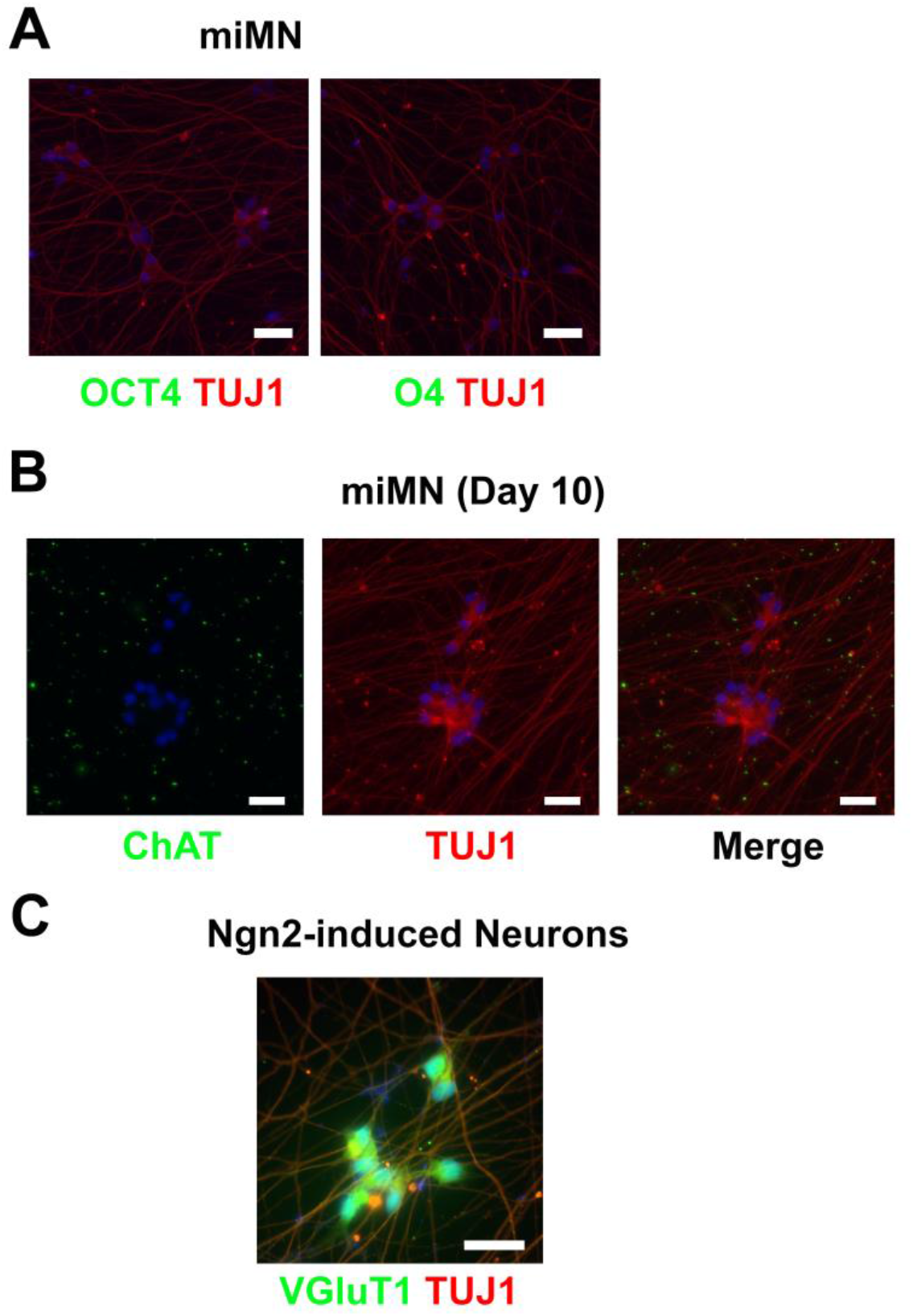
**(A)** TUJ1+ miMNs from the iPSC1 line (day 7 of differentiation) show no detectable expression of the pluripotent stem cell marker (OCT4) and the oligodendrocyte lineage marker (O4). **(B)** TUJ1+ miMNs from the iPSC1 line were immunostained for the cholinergic neuron marker ChAT. Cytoplasmic signal of the ChAT protein was not detected in these miMNs at day 10 of differentiation, compared to cytoplasmic ChAT signal in miMNs at day 20 of differentiation (Figure 3A). **(C)** The iPSC line iPSC1 was differentiated by 3 daily transfections of NSA mRNA alone without using OSA mRNA and SHH/DAPT. TUJ1+ NSA-induced neurons (day 30 of differentiation) express the glutamatergic neuron marker VGlut1, validating the VGluT1 antibody also used in Figure 3E and supporting the requirement of Olig2 for MN induction. Cell nuclei were counterstained with DAPI (Bar = 20 μm).

**Supplemental Figure 2.**
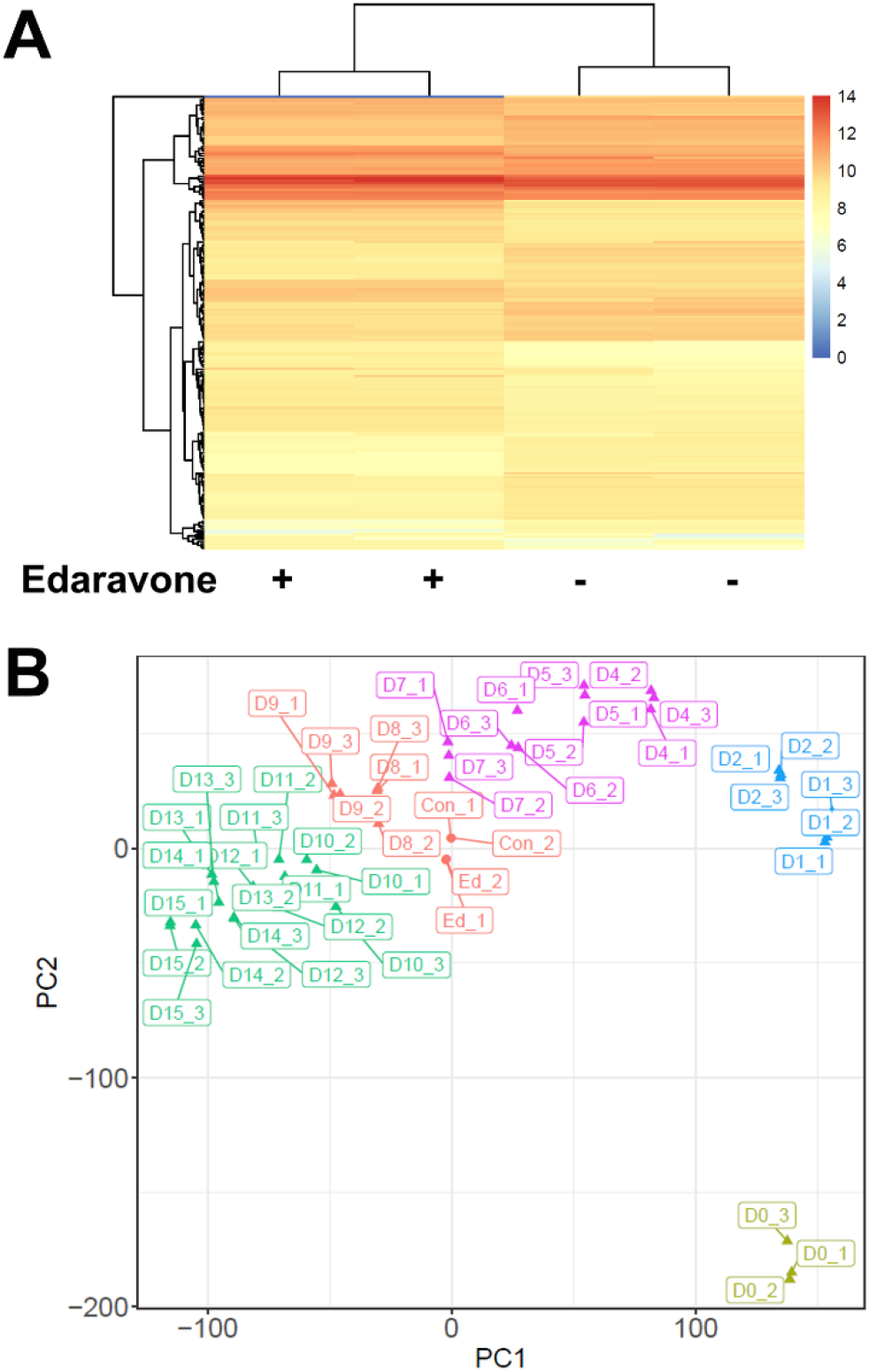
**(A)** Reproducibility of two replicates from the transcriptomic analysis of miMNs with +/- edaravone treatment. Heatmap clustering of RNA-Seq results from miMNs with +/- edaravone treatment (10 µM, 24h, n=2 for each group). Gene expression was calculated by reads per kilobase of transcript, per million mapped reads (RPKM). **(B)** Transcriptomic comparison between miMNs and differentiating cells at various days during compound-induced MN differentiation from human ESCs (Reference 39, GSE140747 from the GEO database). PCA plotting shows that both control (Con_1, Con_2) and edaravone-treated (Ed_1 and Ed_2) miMN samples (two technical replicates) more closely cluster with compound-induced MNs at day 8 and 9 of differentiation (three technical replicates labelled as D8_1/D8_2/D8_3 and D9_1/D9_2/D9_3, respectively). All samples are labelled as “day at differentiation (D)_replicate number”. All samples are grouped and colored based on the clustering result using model-based clustering of batch-effect–adjusted RNA-seq data.

## Supplemental Tables

**Table S1. Up-regulated genes in normal miMNs after edaravone treatment.**

**Table S2. Down-regulated genes in normal miMNs after edaravone treatment.**

**Table S3. Expression data of all genes detected by RNA-seq.**

**Table S4. IPA signaling pathways enriched in edaravone up-regulated genes.**

**Table S5. IPA signaling pathways enriched in edaravone down-regulated genes.**

**Table S6. qPCR primers and antibodies.**

## References

1. Qiang L FR, Abeliovich A.: Remodeling neurodegeneration: somatic cell reprogramming-based models of adult neurological disorders. Neuron 2013, 78:957–969.

2. Ito D OH, Suzuki N.: Accelerating progress in induced pluripotent stem cell research for neurological diseases. Ann Neurol 2012, 72:167–174.

3. Jung YW HE, Kim KY, et al. : Human induced pluripotent stem cells and neurodegenerative disease: prospects for novel therapies. Curr Opin Neurol 2012, 25:125–130.

4. Brown RH, Al-Chalabi A: Amyotrophic Lateral Sclerosis. The New England journal of medicine 2017, 377:162–172.

5. Mejzini R, Flynn LL, Pitout IL, Fletcher S, Wilton SD, Akkari PA: ALS Genetics, Mechanisms, and Therapeutics: Where Are We Now? Frontiers in neuroscience 2019, 13:1310.

6. Hinchcliffe M, Smith A: Riluzole: real-world evidence supports significant extension of median survival times in patients with amyotrophic lateral sclerosis. Degener Neurol Neuromuscul Dis 2017, 7:61–70.

7. 7. Amyotrophic Lateral Sclerosis/Riluzole Study Group II ea: Dose-ranging study of riluzole in amyotrophic lateral sclerosis. The Lancet 1996, 347:1425–1431.

8. Miller RG MJ, Moore DH: Riluzole for amyotrophic lateral sclerosis (ALS) motor neuron disease (MND). Cochrane Database Syst Rev 2012, 3:1–34.

9. Abe K, Aoki M, al. STe: Safety and efcacy of edaravone in well defned patients with amyotrophic lateral sclerosis: a randomised, double-blind, placebo-controlled trial. The Lancet Neurology 2017, 16:505–512.

10. 10. Turnbull J: Reappraisal of an ALS trial: unaccounted procedural risk. The Lancet Neurology 2020, 19:717–718.

11. Mora JS: Edaravone for treatment of early-stage ALS. The Lancet Neurology 2017, 16:772.

12. Akimoto M, Nakamura K, Writing Group on behalf of the Edaravone ALSSG: Edaravone for treatment of early-stage ALS - Authors’ reply. The Lancet Neurology 2017, 16:772.

13. Takei K, Watanabe K, Yuki S, Akimoto M, Sakata T, Palumbo J: Edaravone and its clinical development for amyotrophic lateral sclerosis. Amyotroph Lateral Scler Frontotemporal Degener 2017, 18:5–10.

14. Rothstein JD: Edaravone: A New Drug Approved for ALS. Cell 2017, 171:742.

15. Maury Y, Come J, Piskorowski RA, Salah-Mohellibi N, Chevaleyre V, Peschanski M, Martinat C, Nedelec S: Combinatorial analysis of developmental cues efficiently converts human pluripotent stem cells into multiple neuronal subtypes. Nat Biotechnol 2015, 33:89–96.

16. Du ZW, Chen H, Liu H, Lu J, Qian K, Huang CL, Zhong X, Fan F, Zhang SC: Generation and expansion of highly pure motor neuron progenitors from human pluripotent stem cells. Nat Commun 2015, 6:6626.

17. Mizuguchi R, Sugimori M, Takebayashi H, Kosako H, Nagao M, Yoshida S, Nabeshima Y, Shimamura K, Nakafuku M: Combinatorial roles of olig2 and neurogenin2 in the coordinated induction of pan-neuronal and subtype-specific properties of motoneurons. Neuron 2001, 31:757–771.

18. Mazzoni EO, Mahony S, Closser M, Morrison CA, Nedelec S, Williams DJ, An D, Gifford DK, Wichterle H: Synergistic binding of transcription factors to cell-specific enhancers programs motor neuron identity. Nature neuroscience 2013, 16:1219–1227.

19. Xue Y, Zhan X, Sun S, Karuppagounder SS, Xia S, Dawson VL, Dawson TM, Laterra J, Zhang J, Ying M: Synthetic mRNAs Drive Highly Efficient iPS Cell Differentiation to Dopaminergic Neurons. Stem Cells Transl Med 2019, 8:112–123.

20. Sagal J, Zhan X, Xu J, Tilghman J, Karuppagounder SS, Chen L, Dawson VL, Dawson TM, Laterra J, Ying M: Proneural transcription factor Atoh1 drives highly efficient differentiation of human pluripotent stem cells into dopaminergic neurons. Stem Cells Transl Med 2014, 3:888–898.

21. De Filippis L, Lamorte G, Snyder EY, Malgaroli A, Vescovi AL: A novel, immortal, and multipotent human neural stem cell line generating functional neurons and oligodendrocytes. Stem Cells 2007, 25:2312–2321.

22. Kim D, Paggi JM, Park C, Bennett C, Salzberg SL: Graph-based genome alignment and genotyping with HISAT2 and HISAT-genotype. Nat Biotechnol 2019, 37:907–915.

23. Pertea M, Pertea GM, Antonescu CM, Chang TC, Mendell JT, Salzberg SL: StringTie enables improved reconstruction of a transcriptome from RNA-seq reads. Nat Biotechnol 2015, 33:290–295.

24. Robinson MD, McCarthy DJ, Smyth GK: edgeR: a Bioconductor package for differential expression analysis of digital gene expression data. Bioinformatics 2010, 26:139–140.

25. Robinson MD, Oshlack A: A scaling normalization method for differential expression analysis of RNA-seq data. Genome Biol 2010, 11:R25.

26. Ohta Y, Nomura E, Shang J, Feng T, Huang Y, Liu X, Shi X, Nakano Y, Hishikawa N, Sato K, et al: Enhanced oxidative stress and the treatment by edaravone in mice model of amyotrophic lateral sclerosis. J Neurosci Res 2019, 97:607–619.

27. Richner M, Jager SB, Siupka P, Vaegter CB: Hydraulic Extrusion of the Spinal Cord and Isolation of Dorsal Root Ganglia in Rodents. Journal of visualized experiments : JoVE 2017.

28. Tanabe Y RH, Jessell TM: Induction of motor neurons by Sonic hedgehog is independent of floor plate differentiation. Curr Biol 1995, 5:651–658.

29. 29. Roelink TQCH: The Notch response inhibitor DAPT enhances neuronal differentiation in embryonic stem cell-derived embryoid bodies independently of sonic hedgehog signaling. Dev Dyn 2007, 236:886–892.

30. Zhang Y, Pak C, Han Y, Ahlenius H, Zhang Z, Chanda S, Marro S, Patzke C, Acuna C, Covy J, et al: Rapid single-step induction of functional neurons from human pluripotent stem cells. Neuron 2013, 78:785–798.

31. Wang C, Ward ME, Chen R, Liu K, Tracy TE, Chen X, Xie M, Sohn PD, Ludwig C, Meyer-Franke A, et al: Scalable Production of iPSC-Derived Human Neurons to Identify Tau-Lowering Compounds by High-Content Screening. Stem cell reports 2017, 9:1221–1233.

32. Akopian AN, Sivilotti L, Wood JN: A tetrodotoxin-resistant voltage-gated sodium channel expressed by sensory neurons. Nature 1996, 379:257–262.

33. Khodakhah K, Melishchuk A, Armstrong CM: Killing K channels with TEA+. Proc Natl Acad Sci U S A 1997, 94:13335–13338.

34. Sances S, Bruijn LI, Chandran S, Eggan K, Ho R, Klim JR, Livesey MR, Lowry E, Macklis JD, Rushton D, et al: Modeling ALS with motor neurons derived from human induced pluripotent stem cells. Nat Neurosci 2016, 19:542–553.

35. Johnson MA, Weick JP, Pearce RA, Zhang SC: Functional neural development from human embryonic stem cells: accelerated synaptic activity via astrocyte coculture. J Neurosci 2007, 27:3069–3077.

36. Takazawa T, Croft GF, Amoroso MW, Studer L, Wichterle H, Macdermott AB: Maturation of spinal motor neurons derived from human embryonic stem cells. PLoS One 2012, 7:e40154.

37. Wainger BJ, Kiskinis E, Mellin C, Wiskow O, Han SS, Sandoe J, Perez NP, Williams LA, Lee S, Boulting G, et al: Intrinsic membrane hyperexcitability of amyotrophic lateral sclerosis patient-derived motor neurons. Cell reports 2014, 7:1–11.

38. Michela Dell’Orco PM, Cristina Cereda: Hydrogen Peroxide-Mediated Induction of SOD1 Gene Transcription Is Independent From Nrf2 in a Cellular Model of Neurodegeneration. Biochim Biophys Acta 2015, 1859:315–323.

39. Rayon T, Stamataki D, Perez-Carrasco R, Garcia-Perez L, Barrington C, Melchionda M, Exelby K, Lazaro J, Tybulewicz VLJ, Fisher EMC, Briscoe J: Species-specific pace of development is associated with differences in protein stability. Science 2020, 369.

40. Mammen PP, Shelton JM, Ye Q, Kanatous SB, McGrath AJ, Richardson JA, Garry DJ: Cytoglobin is a stress-responsive hemoprotein expressed in the developing and adult brain. The journal of histochemistry and cytochemistry : official journal of the Histochemistry Society 2006, 54:1349–1361.

41. Zhu J, Garcia-Barcelo MM, Tam PK, Lui VC: HOXB5 cooperates with NKX2-1 in the transcription of human RET. PLoS One 2011, 6:e20815.

42. Leon TY, Ngan ES, Poon HC, So MT, Lui VC, Tam PK, Garcia-Barcelo MM: Transcriptional regulation of RET by Nkx2-1, Phox2b, Sox10, and Pax3. Journal of pediatric surgery 2009, 44:1904–1912.

43. Airaksinen MS, Saarma M: The GDNF family: signalling, biological functions and therapeutic value. Nat Rev Neurosci 2002, 3:383–394.

44. Warren L, Manos PD, Ahfeldt T, Loh YH, Li H, Lau F, Ebina W, Mandal PK, Smith ZD, Meissner A, et al: Highly efficient reprogramming to pluripotency and directed differentiation of human cells with synthetic modified mRNA. Cell Stem Cell 2010, 7:618–630.

45. Schlaeger TM, Daheron L, Brickler TR, Entwisle S, Chan K, Cianci A, DeVine A, Ettenger A, Fitzgerald K, Godfrey M, et al: A comparison of non-integrating reprogramming methods. Nat Biotechnol 2015, 33:58–63.

46. Poleganov MA, Eminli S, Beissert T, Herz S, Moon JI, Goldmann J, Beyer A, Heck R, Burkhart I, Barea Roldan D, et al: Efficient Reprogramming of Human Fibroblasts and Blood-Derived Endothelial Progenitor Cells Using Nonmodified RNA for Reprogramming and Immune Evasion. Hum Gene Ther 2015, 26:751–766.

47. Goparaju SK, Kohda K, Ibata K, Soma A, Nakatake Y, Akiyama T, Wakabayashi S, Matsushita M, Sakota M, Kimura H, et al: Rapid differentiation of human pluripotent stem cells into functional neurons by mRNAs encoding transcription factors. Scientific reports 2017, 7:42367.

48. Serena Stanga LB, Pascal Kienlen-Campard: A Role for GDNF and Soluble APP as Biomarkers of Amyotrophic Lateral Sclerosis Pathophysiology. Front Neurol 2018; 9: 384 2018, 9:384.

49. LiJun Wang YL, Imaharu Nakano: Neuroprotective effects of glial cell line-derived neurotrophic factor mediated by an adeno-associated virus vector in a transgenic animal model of amyotrophic lateral sclerosis. J Neurosci 2002, 22:6920–6928.

50. Thomsen GM, Avalos P, Ma AA, Alkaslasi M, Cho N, Wyss L, Vit JP, Godoy M, Suezaki P, Shelest O, et al: Transplantation of Neural Progenitor Cells Expressing Glial Cell Line-Derived Neurotrophic Factor into the Motor Cortex as a Strategy to Treat Amyotrophic Lateral Sclerosis. Stem Cells 2018, 36:1122–1131.

51. Zhong Zhao DJL, Lap Ho, Sara Bonini, Belinda Shao, Stephen R Salton, Sunil Thomas, and Giulio Maria Pasinetti: Vgf Is a Novel Biomarker Associated With Muscle Weakness in Amyotrophic Lateral Sclerosis (ALS), With a Potential Role in Disease Pathogenesis. Int J Med Sci 2008, 5:92–99.

52. Y. Noda SM, S. Nakamura, M. Shimazawa and H. Hara Neuropeptide VGF-Derived Peptide LQEQ-19 has Neuroprotective Effects in an In Vitro Model of Amyotrophic Lateral Sclerosis. Neurochemical Research 2019, 44:897–904.

53. Masamitsu Shimazawa HT, Hideaki Hara, et al: An Inducer of VGF Protects Cells against ER Stress-Induced Cell Death and Prolongs Survival in the Mutant SOD1 Animal Models of Familial ALS. PLoS One 2010, 5:e15307.

54. Mandolesi G, Gargano S, Pennuto M, Illi B, Molfetta R, Soucek L, Mosca L, Levi A, Jucker R, Nasi S: NGF-dependent and tissue-specific transcription of vgf is regulated by a CREB-p300 and bHLH factor interaction. FEBS Lett 2002, 510:50–56.

55. Alder J, Thakker-Varia S, Bangasser DA, Kuroiwa M, Plummer MR, Shors TJ, Black IB: Brain-derived neurotrophic factor-induced gene expression reveals novel actions of VGF in hippocampal synaptic plasticity. J Neurosci 2003, 23:10800–10808.

56. Cerchia L, D’Alessio A, Amabile G, Duconge F, Pestourie C, Tavitian B, Libri D, de Franciscis V: An autocrine loop involving ret and glial cell-derived neurotrophic factor mediates retinoic acid-induced neuroblastoma cell differentiation. Mol Cancer Res 2006, 4:481–488.

57. Cocco C, Corda G, Lisci C, Noli B, Carta M, Brancia C, Manca E, Masala C, Marrosu F, Solla P, et al: VGF peptides as novel biomarkers in Parkinson’s disease. Cell Tissue Res 2020, 379:93–107.

58. Leigh PN, Meldrum BS: Excitotoxicity in ALS. Neurology 1996, 47:S221–227.

